# Inositol Polyphosphate-4-Phosphatase Type II promotes gemcitabine resistance in pancreatic ductal adenocarcinoma cells via lysosomal exocytosis

**DOI:** 10.64898/2026.08.21.746312

**Authors:** Ché MP Melo, Candaice Newell, Golam Tanjib Saffi, Nathan Ng, Cuyler Yu, Cheng An Wang, Lydia To, Jonathan Tak-Sum Chow, Leonardo Salmena

## Abstract

Chemotherapy resistance is a major challenge in pancreatic ductal adenocarcinoma (PDAC). While high Inositol Polyphosphate-4-Phosphatase Type II (INPP4B) expression correlates with poor outcomes, its function in chemotherapy response is unclear. We show that INPP4B promotes gemcitabine resistance by enhancing lysosomal exocytosis. Across PDAC models, high INPP4B linked to reduced gemcitabine sensitivity, while knockdown restored it. INPP4B also conferred cross-resistance to agents including irinotecan, oxaliplatin, paclitaxel, and daunorubicin. Mechanistically, INPP4B increased cell-surface LAMP1, enhanced extracellular gemcitabine release, and mitigated DNA damage. Pharmacological targeting of lysosomes with chloroquine (CQ), Bafilomycin A (BafA), or specific PIKfyve or TRPML1 inhibitors blocked exocytosis and reversed resistance *in vitro*. Moreover, chloroquine co-treatment restored gemcitabine sensitivity in INPP4B-overexpressing xenografts. These results establish INPP4B-driven lysosomal exocytosis as a key mechanism of gemcitabine resistance, highlighting a therapeutic target for PDAC resensitization.

## INTRODUCTION

Pancreatic ductal adenocarcinoma (PDAC) is among the most aggressive and lethal solid malignancies ^1^. Despite the implementation of aggressive treatment modalities including surgical resection, radiotherapy, chemotherapy, and targeted therapeutic agents, clinical outcomes remain poor ^2^. Therapeutic failure in PDAC is attributed to asymptomatic disease presentation in early stages, early metastatic potential, inherent tumor heterogeneity, an immunosuppressive tumor microenvironment, and substantial chemotherapeutic resistance ^3–5^. Projections indicate that, within the next two decades, PDAC may surpass colorectal cancer, to join lung cancer in becoming the leading causes of cancer-related deaths ^6–8^.

In PDAC, gemcitabine continues to serve as a cornerstone first-line agent in unresectable or metastatic settings, typically administered as monotherapy or widely used in combination with Nab-paclitaxel or capecitabine ^9–11^. Gemcitabine demonstrates broad-spectrum anticancer activity across multiple malignancies, including non-small cell lung carcinoma, breast, ovarian, and bladder cancers ^12^. As a pyrimidine antimetabolite, gemcitabine exerts its cytotoxic effects by inducing S-phase arrest, disrupting DNA synthesis, and triggering apoptosis ^13^. Despite demonstrated efficacy in other malignancies, intrinsic and acquired resistance to gemcitabine remains prevalent among PDAC patients ^13,14^. Resistance in PDAC arises through diverse mechanisms, including impaired nucleoside transport, dysregulated drug metabolism, aberrant activation of cellular signaling pathways, stromal remodeling, hypoxic conditions, genomic instability, and metabolic and immune adaptations ^15^. Therefore, understanding PDAC biology is essential to developing strategies that overcome chemotherapy resistance.

To address this challenge, we investigated a gene of interest in our laboratory with emerging roles in PDAC pathogenesis called Inositol Polyphosphate-4-Phosphatase Type II (INPP4B), a lipid phosphatase INPP4B is a phosphatidylinositol (PtdIns) phosphatase that catalyzes the dephosphorylation of PtdIns(3,4)P₂ to produce PtdIns(3)P at plasma membrane and endosomal membranes ^16–19^. We identified, and subsequent studies have corroborated, a significant association between elevated *INPP4B* expression and poor clinical outcomes in PDAC ^20–24^. Our more recent work demonstrated that INPP4B promotes aggressive PDAC phenotypes by regulating lysosomal function and dynamics ^25,26^. Specifically, INPP4B overexpression enhances lysosomal exocytosis through activation of an INPP4B→PIKfyve→TRPML-1 signalling axis, promoting PDAC cell growth, migration, and invasion ^25,26^. Emerging evidence points to elevated INPP4B expression as a critical mediator of gemcitabine resistance, positioning it as both a prognostic biomarker and a therapeutic target in PDAC ^27–33^.

Lysosomes are membrane-bound organelles essential for cellular degradation processes ^34^. Beyond their traditional catabolic functions, lysosomes serve critical roles in supporting cancer cell growth, survival, and adaptation ^35,36^. Through their ability to coordinate oncogenic signalling pathways, regulate metabolic adaptation, manage cellular stress, and facilitate invasion and metastasis, lysosomes are emerging as key regulators of cancer progression ^37,38^. Crucially, a growing body of evidence shows that lysosomes drive chemoresistance by sequestering lipophilic weak-base anticancer agents including anthracyclines, imidazoacridinones, sunitinib, and pyrimethamine thereby reducing drug access to intracellular targets and promoting multidrug resistance ^39–46^. Lysosomal exocytosis of drug-loaded vesicles can further enhance resistance by clearing chemotherapeutics from cancer cells ^46^. In PDAC, lysosomes support therapy resistance through drug sequestration, autophagy-dependent survival, and stress-adaptive metabolic recycling; accordingly, chloroquine-mediated lysosome inhibition sensitizes PDAC cells to replication stress by disrupting nucleotide biosynthesis and depleting aspartate ^47^.

Although INPP4B is a reported prognostic marker and driver of PDAC aggressiveness, its contribution to treatment outcomes has not been defined. We hypothesized that INPP4B drives gemcitabine resistance, providing a mechanistic explanation for the poor prognosis associated with its elevated expression. We show that INPP4B drives gemcitabine resistance by promoting lysosomal exocytosis, which enhances drug efflux and decreases intracellular retention. This resistance is reversible, as pharmacologically inhibiting lysosomes restored drug sensitivity *in vitro* and re-sensitized tumors to gemcitabine *in vivo*. Additionally, this mechanism confers resistance to various chemotherapeutic classes, establishing lysosomal exocytosis as a putative key therapeutic target for overcoming chemoresistance in PDAC.

## RESULTS

### Inpp4b drives proliferation, invasion, and tumor growth in murine PDAC models

INPP4B overexpression can drive growth and proliferative phenotypes in human PDAC cell lines ^25,26^. To test whether these phenotypes are conserved in a primary murine setting, Inpp4b-overexpressing and knockdown PDAC cell lines were generated from tumor cells derived from *LSL-Kras^G12D/+^; LSL-Trp53^R172H/+^; Pdx1-Cre* (KPC) mice. Murine Inpp4b was stably overexpressed in KPC cells *via* lentiviral transduction with the *pSMAL* vector ^48^ to generate *KPC-Inpp4b* and *KPC-pSMAL* control cells (**Figure 1a**). *Inpp4b* was knocked down by lentiviral transduction with pLKO.1 shRNAs targeting *Inpp4b* (KPC-sh36, KPC-sh38), alongside a non-targeting control (KPC-NTC)(**Figure 1b**). Consistent with previous findings, Inpp4b overexpression increased cell growth (**Figure 1c**), while knockdown reduced cell growth (**Figure 1d**) ^25^. In 3D spheroid assays KPC-*Inpp4b* cells formed larger, more invasive spheroids compared to KPC-*pSMAL* controls (**Figure 1e,f**). These findings show that Inpp4b promotes proliferative and invasive phenotypes in a primary murine PDAC model, confirming that its growth-promoting roles are conserved across human and mouse pancreatic cancer systems.

**Figure 1:**
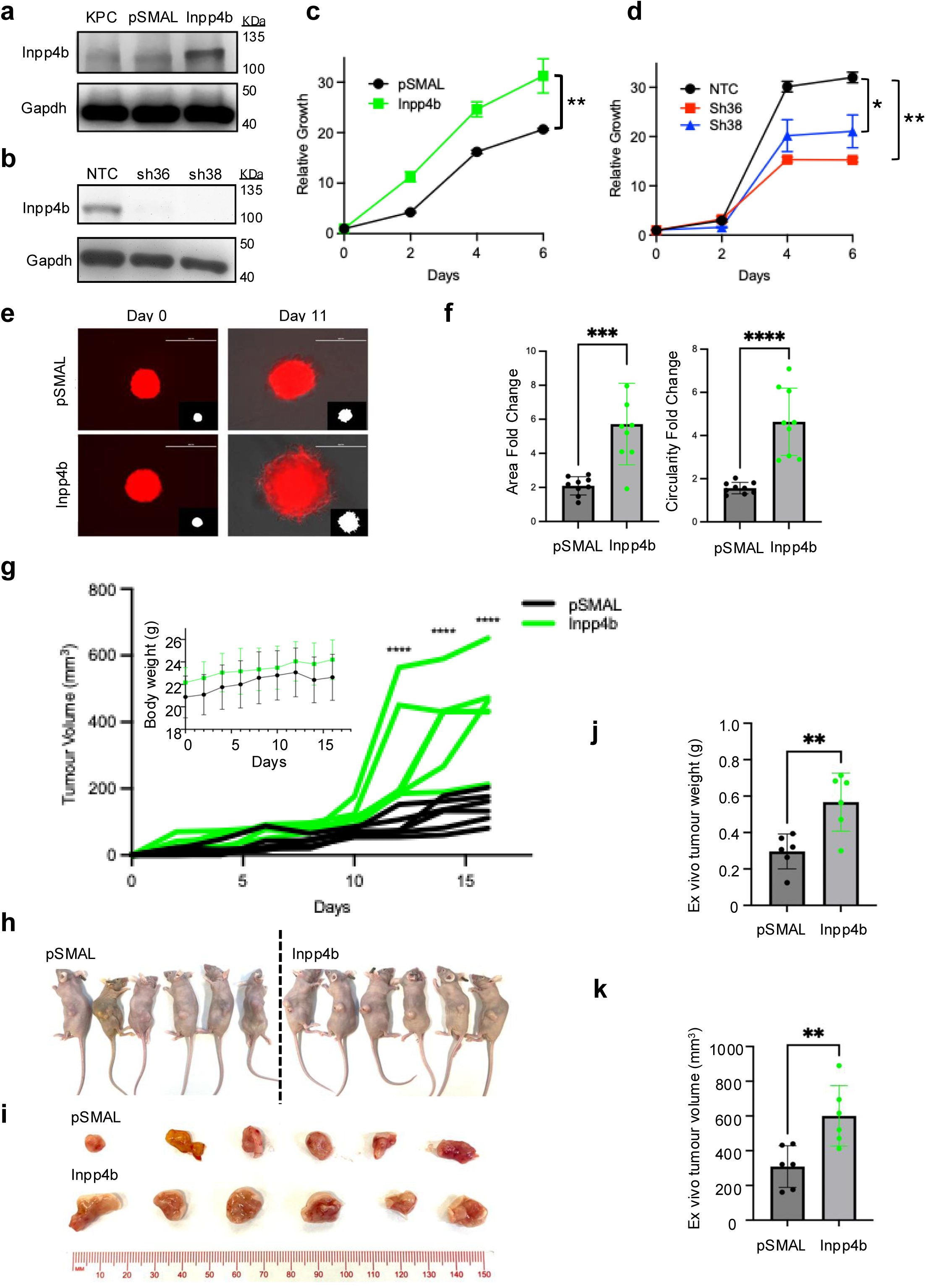
Inpp4b regulates KPC cell proliferation, invasive capacity, and tumorigenic potential. Western blot validation of KPC cell lines following INPP4B overexpression (**a**) and shRNA knockdown (**b**). Growth analysis of Inpp4b-overexpressing (**c**) and knockdown (**d**) KPC cells. (e) Representative images and masks of Inpp4b-overexpressing 3D spheroids (red fluorescence, scale bar 400 µm). (**f**) Fold change in spheroid area (left) and circularity (right). (**g**) Mouse body weight and tumor volume monitored over 16 days. (**h**) Post-mortem tumor specimens and (**i**) dissected tumors grouped by genotype (*pSMAL* vs. *Inpp4b*). Ex-vivo tumor weight (**j**) and volume (**k**). *p < 0.05, **p < 0.01, ***p < 0.001, ****p < 0.0001; ns, not significant. Data represent mean ± SEM from ≥3 independent experiments. Statistical significance determined by one-way ANOVA (**c, d, g**) or unpaired t-test (**f, j, k**).

To extend these findings *in vivo*, *KPC-pSMAL* and *KPC-Inpp4b* cells were engrafted subcutaneously into the flanks of immunodeficient nude mice. In these cell-derived xenograft (CDX) models, KPC-*Inpp4b* tumors grew significantly larger than controls, with no consequences in total body weight (**Figure 1g-k**) further supporting a role for Inpp4b in promoting PDAC tumor aggressiveness.

### INPP4B Levels Impact Gemcitabine Response and Resistance in PDAC Cells

While INPP4B is linked to poor prognosis and tumor aggressiveness in PDAC, its impact on treatment response remains unclear. To evaluate the therapeutic relevance of INPP4B expression in PDAC, we analyzed data from the TCGA-PAAD database in a cohort of 100 gemcitabine-treated patients ^49^. In this cohort, patients with high *INPP4B* expression exhibited a significantly poorer overall survival (**Figure 2a**), suggesting that INPP4B may act as a determinant of gemcitabine resistance in PDAC. To further evaluate if INPP4B affects gemcitabine responsiveness globally, we correlated *INPP4B* transcript expression from the Cancer Cell Line Encyclopedia (CCLE) with gemcitabine response data from the Cancer Therapeutics Response Portal (CTRP). We analyzed three datasets covering CCLE cell lines treated with gemcitabine alone, or in combination with tanespimycin or navitoclax. In all three conditions, *INPP4B* correlated positively with area under the curve (AUC), indicating resistance to gemcitabine in cells with higher *INPP4B* (**Figure 2b and Figure S1a,b)**. Since combination with tanespimycin or navitoclax neither strengthened nor diminished this relationship, the association appears driven specifically by gemcitabine, suggesting a broader role for INPP4B in mediating gemcitabine response. To validate this association experimentally, we measured INPP4B protein expression by western blot across a panel of eight human PDAC cell lines (**Figure 2c**). We then performed gemcitabine dose-response evaluations to determine the IC50 for each cell line (**Figure 2d,e**). Consistently, INPP4B expression correlated positively with gemcitabine IC50 (r = 0.797; *P*-value = 0.018; **Figure 2f**), confirming that gemcitabine resistance is directly correlated with INPP4B expression. These data suggest that high INPP4B expression limits gemcitabine response, potentially explaining the poor clinical outcomes associated with high INPP4B in PDAC patients.

**Figure 2:**
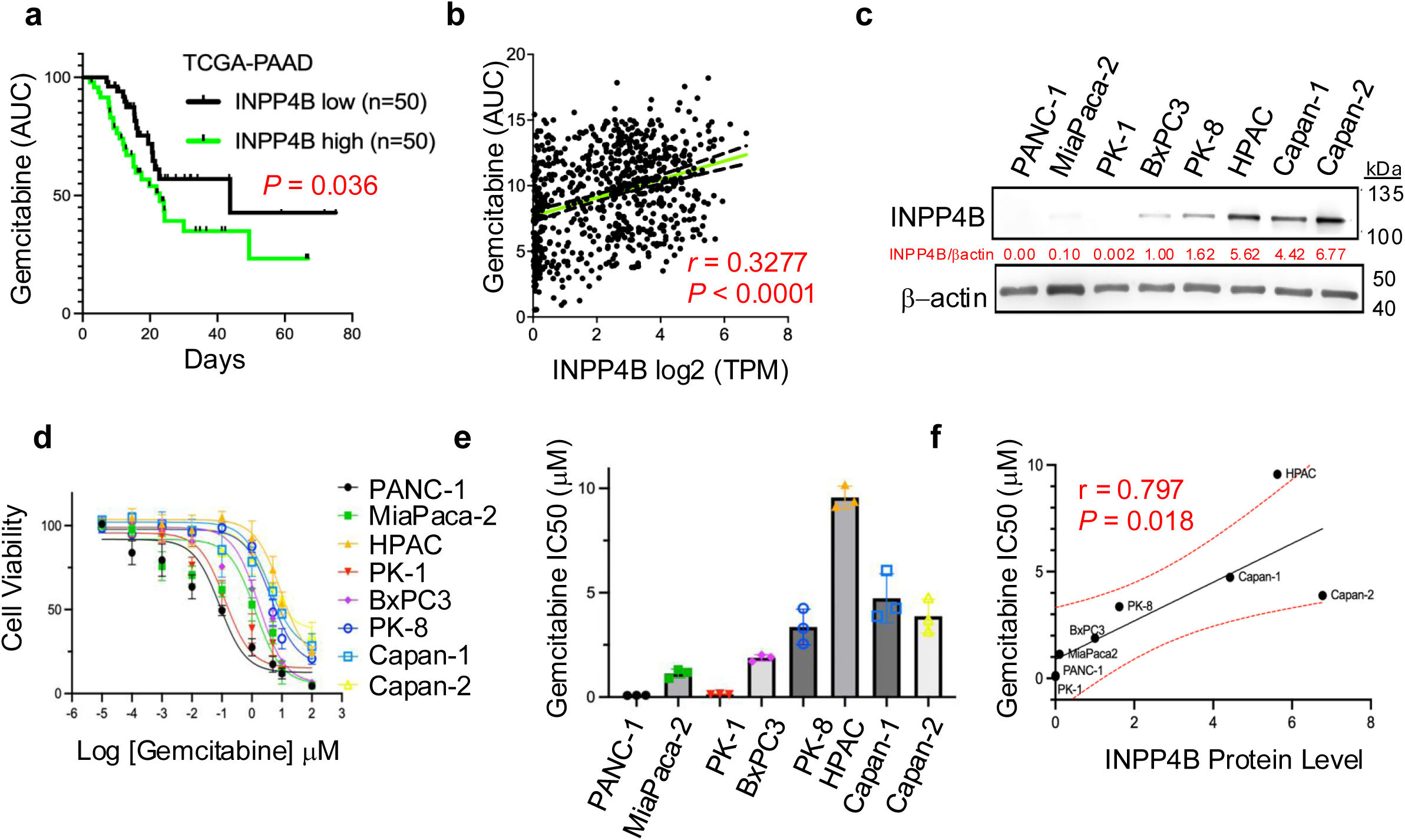
INPP4B expression correlates with gemcitabine chemoresistance in PDAC. **(a)** Kaplan–Meier overall survival (OS) analysis of gemcitabine-treated PDAC patients (TCGA-PAAD) by *INPP4B* expression. **(b)** Correlation between *INPP4B* expression (CCLE) and gemcitabine AUC (CTRP). **(c)** Western blot of INPP4B expression in 8 human PDAC cell lines (β-actin loading control). **(d)** Representative gemcitabine dose–response curves for the cell lines. **(e)** Histogram of g emcitabine IC50 values (µM). **(f)** Pearson correlation between basal INPP4B protein and gemcitabine IC50. *p* < 0.05, \*\**p* < 0.01, \*\*\**p* < 0.001, \*\*\*\**p* < 0.0001; ns, not significant. All experiments were performed in triplicate.

To determine if Inpp4b plays a direct functional role in gemcitabine resistance, we manipulated its expression in KPC cells and assessed gemcitabine sensitivity. We observed that Inpp4b overexpression significantly increased the gemcitabine IC_50_ (**Figure 3a,b**), while *Inpp4b* knockdown decreased it (**Figure 3c,d**), a finding corroborated across human PDAC cell lines (**Figure S1c-h**). Furthermore, to assess whether acquired resistance selects for Inpp4b overexpression, we generated gemcitabine resistant KPC cells via stepwise drug escalation (**Figure 3e**). These resistant cells demonstrated a marked rightward shift in dose-response curves and significantly elevated IC_50_ values compared to parental counterparts (**Figure 3f,g**). Crucially, Western blot confirmed a significant, reproducible upregulation of Inpp4b in these resistant cells (**Figure 3h,i**). Collectively, these data suggest that Inpp4b is a functional driver of gemcitabine resistance that is enriched under therapeutic selection pressure in PDAC.

**Figure 3:**
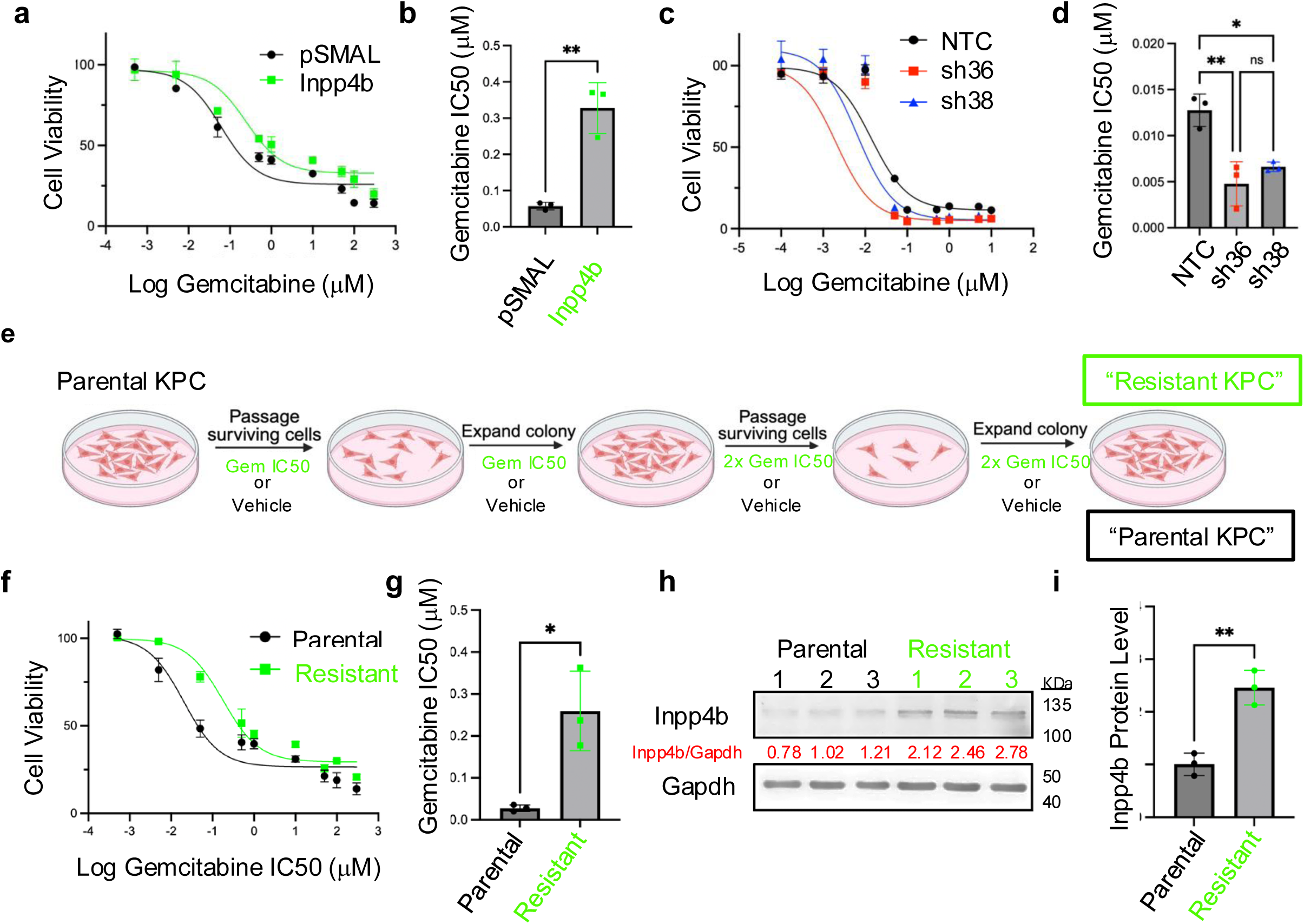
INPP4B drives gemcitabine resistance in PDAC cells. (**a**) Representative gemcitabine DR-curves and **(b)** IC50 values for *KPC-Inpp4b* cells. **(c)** Representative DR-curves and **(d)** IC50 values for KPC knockdown cells. **(e)** Schematic of gemcitabine-resistant KPC cell generation. **(f)** Representative DR-curves and **(g)** comparative IC50 values of gemcitabine-resistant versus parental lines. **(h)** Western blot and **(i)** quantification of INPP4B protein levels in parental and resistant KPC lines (n=3). Significance determined by two-way ANOVA with Sidak’s post hoc (d) or unpaired two-tailed t-test (b, g, i). *p<0.05, **p<0.01, ***p<0.001, ****p<0.0001; ns, not significant.

### Inpp4b-associated gemcitabine resistance is linked to enhanced lysosomal exocytosis

To elucidate the mechanisms by which INPP4B confers gemcitabine resistance, we drew upon our previous research demonstrating that INPP4B increases PDAC cell aggressiveness by promoting lysosomal exocytosis through an INPP4B–PIKfyve–TRPML-1 signaling axis ^25,26^. Building on recent reports linking lysosomal pathways to gemcitabine resistance in PDAC ^50^, we hypothesized that INPP4B exploits this exocytosis pathway to confer a survival advantage under gemcitabine treatment. To investigate this, we evaluated lysosomal exocytosis across our PDAC cell models.

Lysosomal exocytosis is a complex process that culminates in the fusion of lysosomal membranes to the plasma membrane ^51,52^. The process can be quantified by flow cytometry by measuring cell-surface levels of the lysosome-specific protein LAMP1 on live, unfixed, unpermeabilized cells ^25,26^. Using this approach, we assessed baseline cell-surface LAMP1 expression across our panel of human PDAC cell lines (**Figure 4a**). We found a strong positive correlation between cell-surface LAMP1 levels and INPP4B expression across the panel (r = 0.931; *P* = 0.0008; **Figure 4b**), indicating that INPP4B expression is associated with enhanced lysosomal exocytosis in PDAC cells. We next examined the relationship between cell-surface LAMP1 levels and gemcitabine IC_50_ values across the same cell panel and found a similarly strong positive correlation (r = 0.807; *P* = 0.0156; **Figure 4c**). Together, these data point to a putative link between INPP4B expression, lysosomal exocytosis, and gemcitabine resistance.

**Figure 4:**
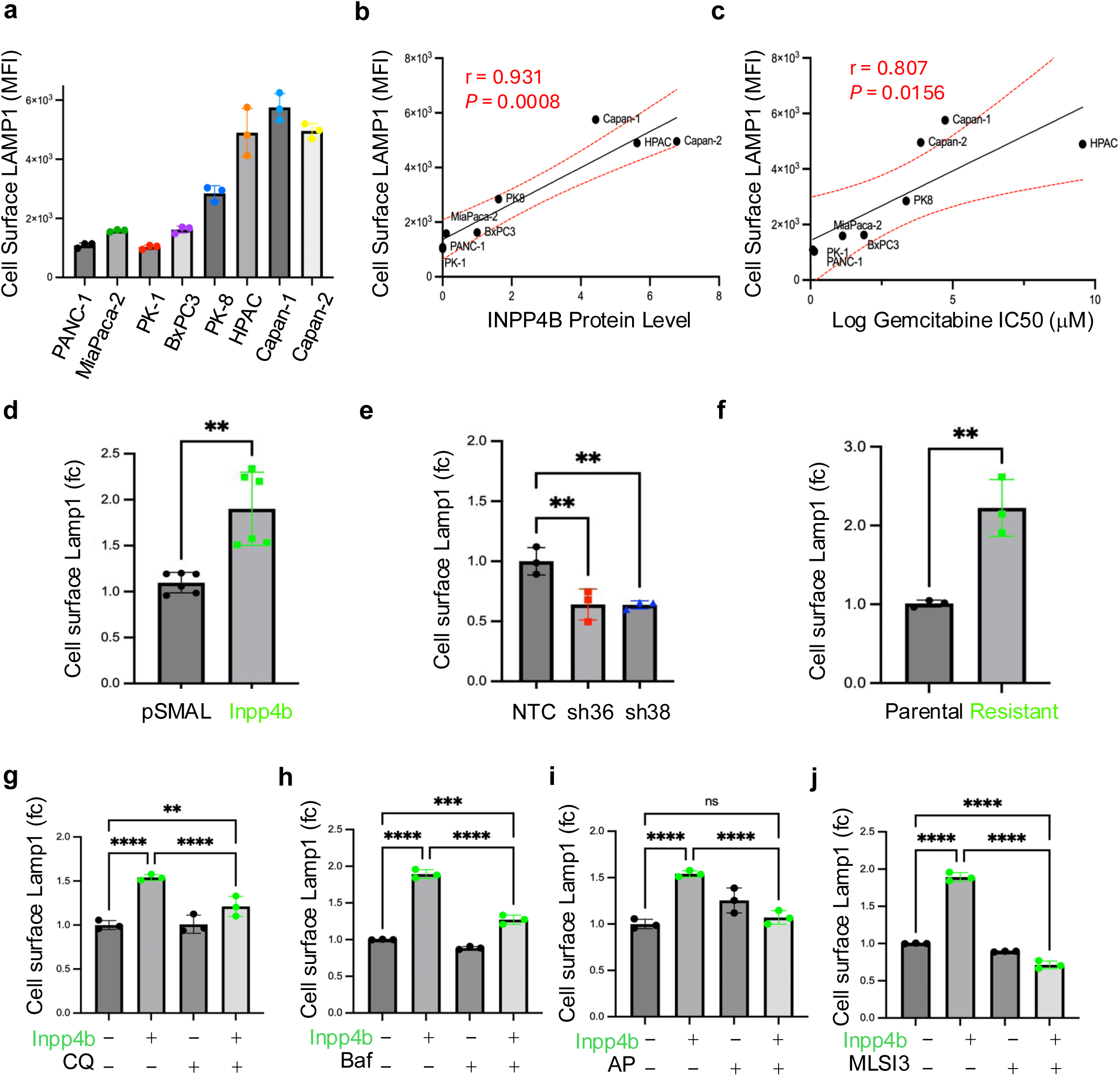
INPP4B regulates LAMP1 surface expression and correlates with gemcitabine response. (**a**) Surface LAMP1 levels in 8 human PDAC cell lines. (**b–c**) Pearson correlation of surface LAMP1 levels with (**b**) INPP4B expression and (**c**) gemcitabine IC50. (**d–f**) Fold change in surface LAMP1 in (**d**) *KPC-Inpp4b* overexpression, (**e**) *KPC-Inpp4b* knockdown, and (**f**) gemcitabine-resistant KPC cells. Cell surface Lamp1 fold change measured by flow-cytometry in KPC control and KPC Inpp4b overexpressing cells following treatment with **(g)** CQ, **(h)** Baf, **(i)** apilimod and **(j)** MLSI3. *p<0.05, **p<0.01, ***p<0.001, ****p<0.0001; ns, not significant. Statistical tests: two-way ANOVA (Sidak’s post hoc) for (a-e) and unpaired t-test for (d, f). All replicate experiments were performed at least three times.

To determine if Inpp4b directly influences exocytosis, we measured cell-surface Lamp1 in KPC gain- and loss-of-function models. KPC-*Inpp4b* cells exhibited nearly a twofold increase in surface Lamp1 compared to KPC-*pSMAL* controls (**Figure 4d**), whereas *Inpp4b* knockdown significantly diminished exocytic activity (**Figure 4e**). Consistent with these results, gemcitabine-resistant KPC cell lines showed a twofold increase in cell-surface Lamp1 relative to vehicle-treated counterparts (**Figure 4f**). Given the link between gemcitabine resistance and INPP4B-driven lysosomal exocytosis, we hypothesized that blocking lysosomal exocytosis would restore drug sensitivity. Lysosomal inhibitors have been shown to modulate exocytosis through their ability to alter organelle pH, calcium homeostasis, and intracellular vesicle trafficking ^53^. To test this in our KPC models, we evaluated four lysosomal inhibitors targeting distinct lysosomal processes: chloroquine (CQ) and bafilomycin A1 (Baf), which inhibit lysosomal acidification; apilimod, a PIKfyve kinase inhibitor; and ML-SI3, a selective TRPML-1 channel antagonist. We next asked whether each lysosome inhibitor could attenuate the elevated exocytosis associated with Inpp4b overexpression. Indeed, Inpp4b overexpression consistently increased exocytosis across KPC cell models (**Figure 4g-j**). While CQ, Baf, apilimod, and ML-SI3 had minimal effect on exocytosis in *KPC-pSMAL* control cells, all four agents significantly reduced the elevated exocytosis observed in *KPC-Inpp4b* cells (**Figure 4g-j**). These results were corroborated in the human PDAC cell line MiaPaCa-2 where cell surface LAMP1 levels were elevated by INPP4B overexpression, and specifically reduced by each of CQ, Baf, AP, and ML-SI3 (**Figure S2a-c)**. These findings demonstrate that pharmacological inhibition of lysosomal function selectively normalizes the elevated exocytic activity driven by Inpp4b overexpression in PDAC models. Collectively, these data suggest that INPP4B-mediated lysosomal exocytosis is a mechanistic driver of gemcitabine resistance in PDAC.

### Lysosomal targeting agents resensitize Inpp4b overexpressing PDAC cells to gemcitabine

To establish baseline sensitivities, we tested the responsiveness of *KPC-pSMAL* and *KPC-Inpp4b* cells to individual lysosomotropic agents, evaluating their effects on viability prior to assessing combination treatments. Dose-response analyses revealed no significant differences in viability between the two cell lines for any of the four compounds (**FigureS3 a–h**), although CQ elicited a dose-dependent cytotoxic response across both genotypes (**FigureS3a,b**). Next, we assessed cell viability across a range of concentration combinations of gemcitabine with CQ, Baf, apilimod, or ML-SI3. *KPC-pSMAL and KPC-Inpp4b* cells were treated in a 10 × 6 matrix of increasing agent combinations, where gemcitabine doses were maintained at sub-IC_50_ doses throughout (**FigureS4 a-d**). Synergy was quantified using the Synergy Finder tool by implementing the Zero Interaction Potency (ZIP) model ^54^ and applying a synergy score threshold of 5, consistent with previous studies ^55–57^.

Synergy analysis of gemcitabine + CQ revealed a slightly antagonistic across-the-board (ATB) interaction in KPC-pSMAL control (ZIP = –5.06) and a below-threshold ATB score in *KPC-Inpp4b* cells (ZIP = 2.87). Despite falling below the synergy threshold overall, *KPC-Inpp4b* cells showed a clear relative shift toward synergy compared with controls (**Figure S5a-c**), with a distinct region of strong synergy identified at 3.0 µM CQ + 37.5 nM gemcitabine that also corresponded to a marked reduction in cell viability (**Figure S5d,e**). A similar pattern emerged for gemcitabine + Baf. *KPC-Inpp4b* cells exhibited a positive ATB ZIP score (8.19) compared with no synergy in KPC-pSMAL controls (ZIP = –0.97) (**Figure S5f-h**). Peak synergy occurred at 5 nM Baf + 4.86 nM gemcitabine, coinciding with reduced viability in *KPC-Inpp4b* cells (**Figure S5i,j**). Gemcitabine + apilimod produced the most pronounced synergy of all combinations tested, with the ATB ZIP score rising to 10.04 in *KPC-Inpp4b* cells versus –2.36 in controls (**Figure S5k-m**). Maximal synergy occurred at 0.04 µM apilimod + 37.5 nM gemcitabine, with a corresponding reduction in *KPC-Inpp4b* cell viability (**Figure S5n,o**). Finally, gemcitabine + ML-SI3 likewise increased the ATB ZIP score in KPC-*Inpp4b* cells (5.36) relative to controls (–4.03) (**Figure S5p-r**), with peak synergy at 1.56 µM ML-SI3 + 18.75 nM gemcitabine and a corresponding decrease in viability (**Figure S5s,t**). In independent experiments measuring cell growth, all four agents consistently reversed Inpp4b-mediated resistance, effectively re-sensitizing *KPC-Inpp4b* cells to gemcitabine (**Figure S6a-d**). In sum, these findings indicate that Inpp4b overexpression confers a selective vulnerability to lysosome-targeting agents when combined with gemcitabine, highlighting a potential therapeutic strategy to overcome Inpp4b-mediated chemoresistance.

### Inpp4b overexpression reduces gemcitabine mediated DNA damage

To corroborate the above findings and determine whether Inpp4b confers resistance by limiting the toxic effects induced by gemcitabine, we measured γH2AX (a marker of DNA double-strand breaks and replication-associated DNA damage) using flow cytometry with a FITC-tagged γH2AX antibody (**Figure S7**). We observed that gemcitabine treatment of *KPC-Inpp4b* cells resulted in significantly lower γH2AX levels compared to *KPC-pSMAL cells* (**Figure 5a,b**). These findings show that Inpp4b attenuates the DNA damage response elicited by gemcitabine, consistent with resistance. Notably, when combined with gemcitabine, all lysosomal inhibitors (CQ, Baf, apilimod, and ML-SI3) enhanced DNA damage in KPC-*Inpp4b* but not in KPC-*pSMAL* cells, where lysosome inhibition had either no effect on or reduced the gemcitabine response (**Figure 5c–j**). By employing DNA damage levels as an intracellular surrogate, our findings provide additional support for the involvement of lysosomal function in gemcitabine resistance. This suggests that enhanced exocytosis mediated by INPP4B could be a pivotal underlying mechanism that impedes gemcitabine-induced DNA damage.

**Figure 5:**
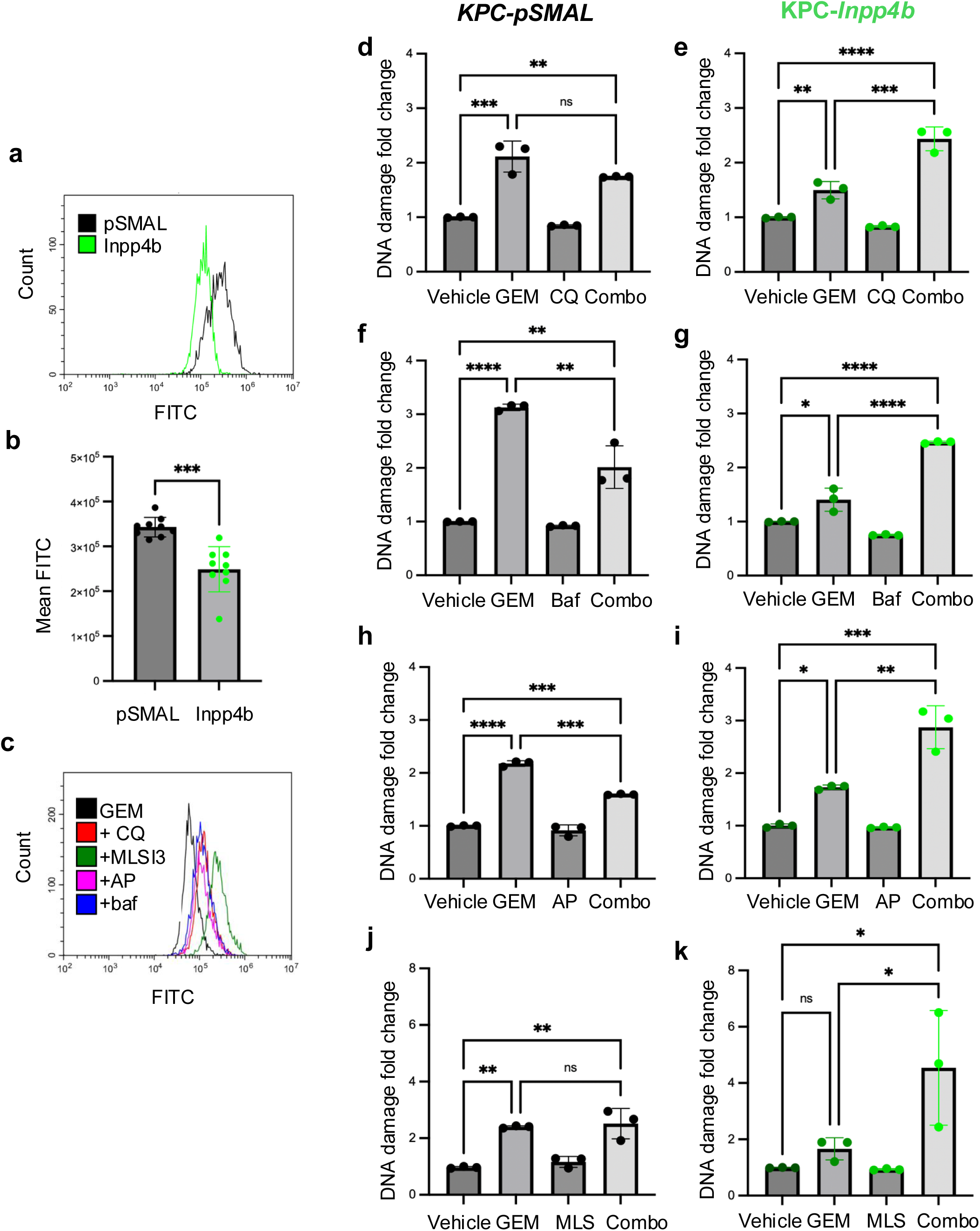
Lysosomal inhibitors enhance gemcitabine-induced DNA damage in an Inpp4b-dependent manner. (**a**) Representative flow cytometry histograms of γH2AX signal (FITC) following 3-hour gemcitabine (GEM) exposure in *KPC-pSMAL* and *KPC-Inpp4b* cells. (**b**) Quantification of gemcitabine-mediated DNA damage (mean FITC intensity, n=9). (**c**) Representative γH2AX histograms in *KPC-Inpp4b* cells exposed to gemcitabine ± CQ, ML-SI3, AP, or Baf. (**d–k**) Fold change in gemcitabine-induced DNA damage following co-treatment with indicated lysosomal inhibitors in *KPC-pSMAL* and *KPC-Inpp4b* cells. Data represent mean ± SEM (n ≥ 3). Significance: *p < 0.05, **p < 0.01, ***p < 0.001, ****p < 0.0001; ns, not significant. Statistical analysis by two-way ANOVA (Sidak’s post hoc) or unpaired t-test (**b**).

### Inpp4b expression promotes gemcitabine exocytosis

To determine whether enhanced exocytosis contributes to Inpp4b-dependent gemcitabine resistance, we quantified extracellular gemcitabine concentrations via liquid chromatography–tandem mass spectrometry (LC–MS/MS; **Figure 6a and Figure S8**). KPC-*Inpp4b* and KPC-*pSMAL* cells were exposed to 10 µM gemcitabine for 3 h, then washed and incubated in drug-free medium. Conditioned media collected sequentially over 90 min revealed significantly higher extracellular gemcitabine in KPC-*Inpp4b* cells compared to KPC-*pSMAL* controls at all time points (**Figure 6b**). To assess if the observed gemcitabine release is lysosome-dependent, gemcitabine treated KPC-*Inpp4b* and KPC-*pSMAL* cells were co-treated with chloroquine (CQ), bafilomycin A1 (Baf), apilimod, or ML-SI3. Lysosomal inhibition did not alter gemcitabine efflux in KPC-*pSMAL* cells, but significantly reduced extracellular gemcitabine accumulation in KPC-*Inpp4b* cells (**Figure 6c**). These findings provide further evidence that Inpp4b overexpression promotes gemcitabine release via lysosomal exocytosis, providing a mechanistic basis for evaluating lysosome-targeting agents as combination therapies.

**Figure 6:**
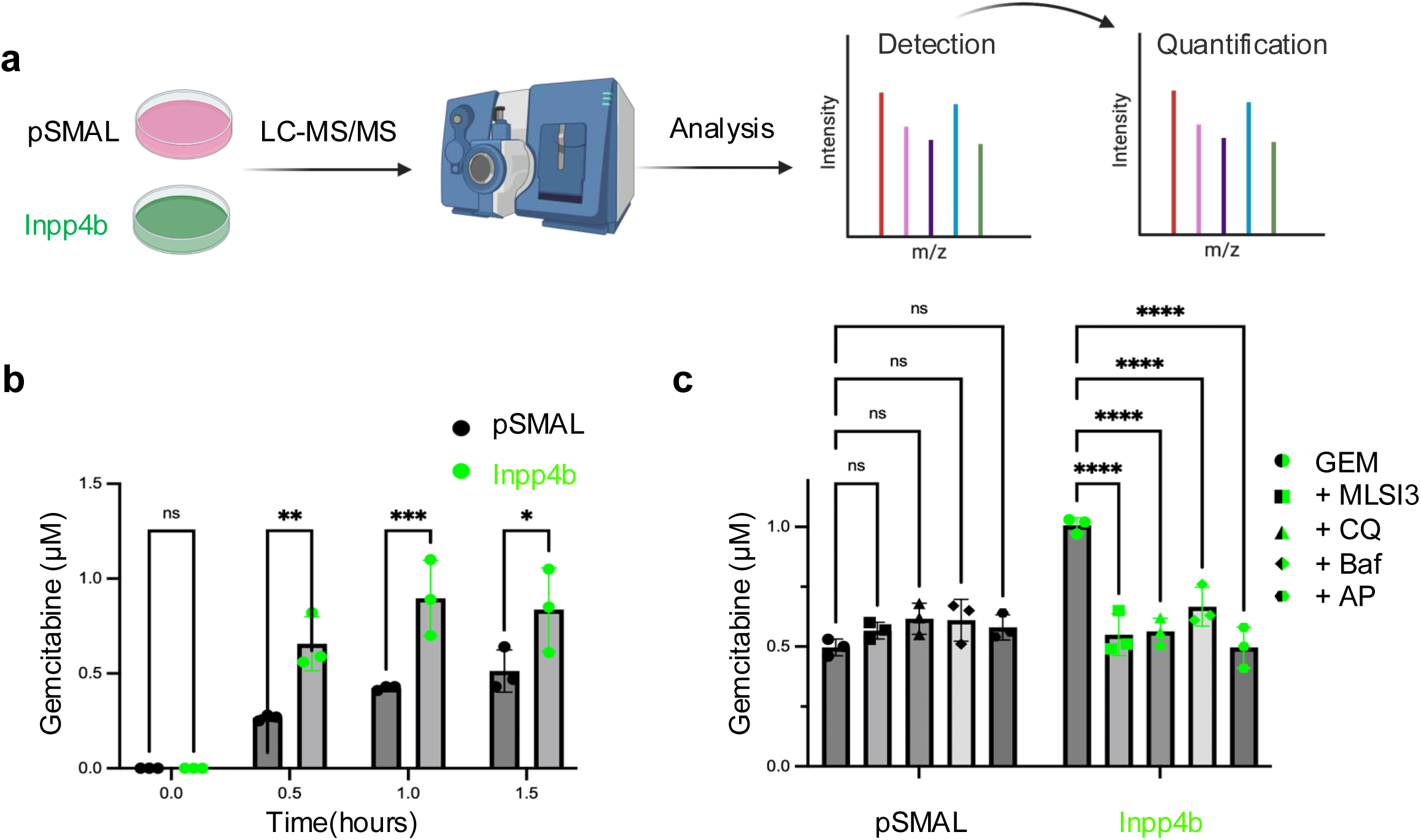
Lysosomal inhibitors modulate extracellular gemcitabine accumulation. **(a)** Schematic of LC–MS/MS workflow. **(b)** Extracellular gemcitabine (µM) quantified via LC–MS/MS over 90 min. **(c)** Extracellular gemcitabine (µM) 1 h post-wash following combination treatment with CQ, Baf, AP, or MLSI3. Data are mean ± SEM (n ≥ 3). Significance determined by one-way ANOVA with Tukey’s post hoc test (*p* < 0.05).

### Inpp4b expression confers multi-drug resistance in PDAC

Given the broad contribution of lysosomes to chemotherapeutic resistance and the established role of INPP4B in regulating lysosomal function, we hypothesized that the Inpp4b-dependent gemcitabine-resistant phenotype in KPC cells may extend to other anticancer agents ^46,58^. We therefore investigated whether *Inpp4b* overexpression confers resistance to mechanistically distinct chemotherapeutic drugs. We evaluated four mechanistically distinct chemotherapeutic classes including irinotecan (a topoisomerase I inhibitor), oxaliplatin (a platinum-based antineoplastic agent), paclitaxel (a microtubule-stabilizing agent), and daunorubicin (an anthracycline), all known to be affected by lysosomal sequestration and exocytosis ^59^. 72-hour dose–response assays revealed that *Inpp4b* overexpression increases resistance to all four agents, as indicated by rightward-shifted dose–response curves and elevated IC50 values (**Figure S9a–b, e–f, i–j, and m–n**). Conversely, Inpp4b knockdown sensitized KPC cells to each agent, reflected by leftward shifts and reduced IC50 values (**Figure S9c–d, g–h, k–l, and o–p**). These findings show that Inpp4b is a mediator of resistance to mechanistically distinct chemotherapeutics. As irinotecan, oxaliplatin, and paclitaxel are clinically relevant in PDAC treatment in the FOLFINNOX regimen, INPP4B-dependent lysosomal regulation may contribute to broader therapeutic resistance and PDAC progression.

### Chloroquine resensitizes Inpp4b overexpression tumours *in vivo*

To validate our findings *in vivo*, KPC-pSMAL control and *KPC-Inpp4b* cells were implanted subcutaneously into the flanks of immunodeficient nude mice. Once tumours reached a mean volume of ≥100 mm³, mice within each cohort were randomized to receive vehicle, gemcitabine (80 mg/kg, once weekly), chloroquine (CQ; 30 mg/kg, daily), or the gemcitabine/CQ combination for two consecutive weeks (**Figure 7a**). Longitudinal tumour-volume measurements showed that, in mice bearing KPC-pSMAL tumours, CQ monotherapy did not significantly affect tumour growth relative to vehicle treatment. In contrast, gemcitabine alone or in combination with CQ significantly reduced tumour volume (**Figure 7b,c and Figure S10a**). As observed previously, *KPC-Inpp4b* tumours exhibited accelerated growth relative to KPC-pSMAL control tumours (**Figure 1g–k**). Notably, gemcitabine monotherapy did not significantly reduce the growth of *KPC-Inpp4b* tumours compared with vehicle-treated controls, consistent with Inpp4b-mediated gemcitabine resistance (**Figure 7d,e and Figure S10b**). However, co-treatment with gemcitabine and CQ restored sensitivity to gemcitabine, resulting in marked suppression of *KPC-Inpp4b* tumour growth to levels comparable to those observed in gemcitabine-treated KPC-pSMAL tumours (**Figure 7d,e and Figure S10b**). Analysis of the tumour inhibition rate (TIR) revealed no significant difference between gemcitabine monotherapy and combination treatment in KPC-pSMAL tumours, whereas the TIR was significantly greater in the combination-treatment group than in the gemcitabine-alone group **Figure S10c**). Together, these findings provide *in vivo* evidence that lysosomal inhibition by CQ overcomes Inpp4b-driven gemcitabine resistance and restores gemcitabine responsiveness in *KPC-Inpp4b* tumours.

**Figure 7:**
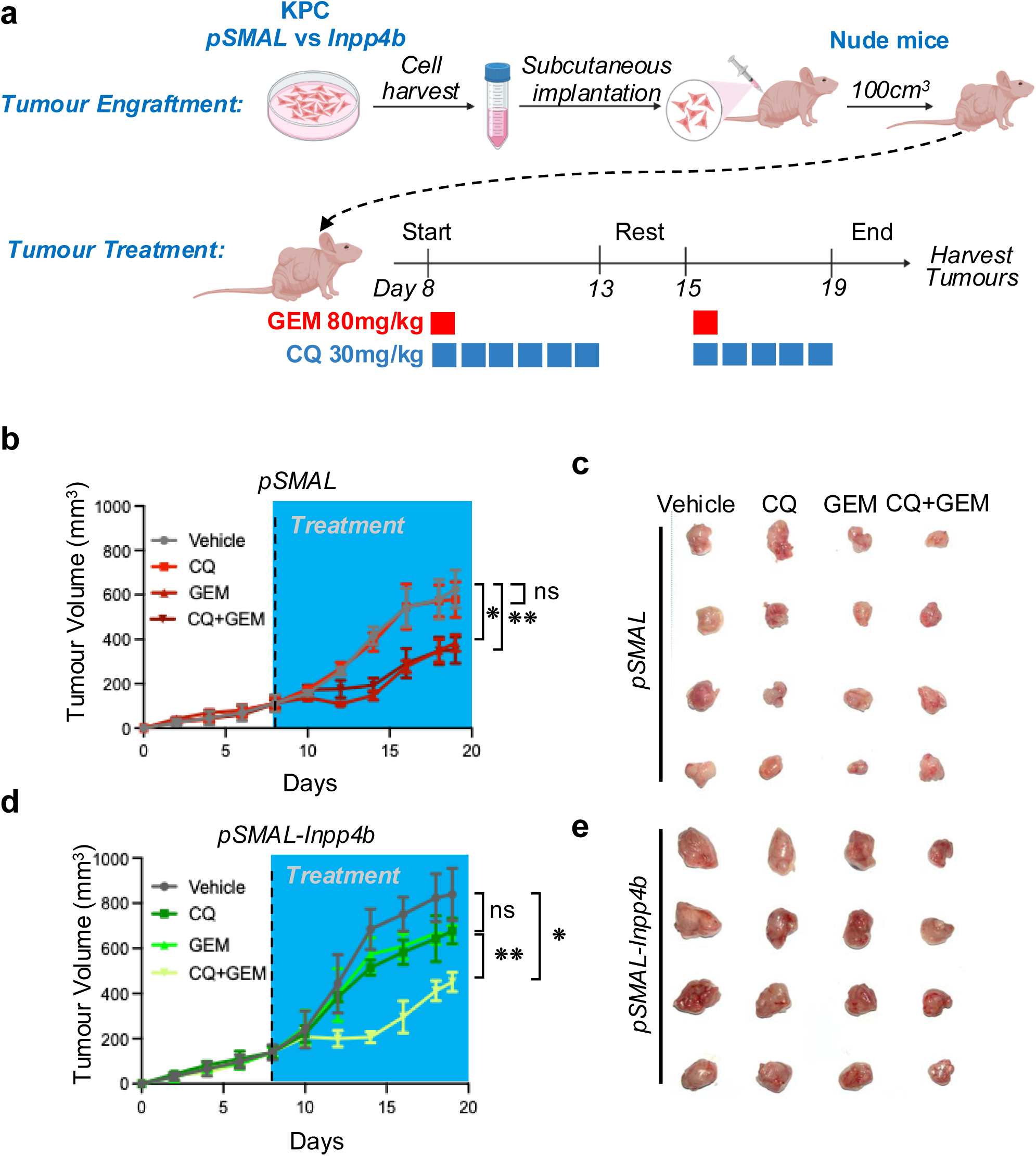
CDX model tumor growth and treatment response in KPC-derived xenografts. (**a**) Experimental workflow timeline. Tumor volume (mm³) monitored over 19 days (blue shading: treatment interval) for (**b**) KPC-*pSMAL* and (**d**) KPC-*Inpp4b* xenografts, with (**c, e**) representative images of excised tumors. Significance: \**p* < 0.05, \*\**p* < 0.01, \*\*\**p* < 0.001, \*\*\*\**p* < 0.0001; ns, not significant. (n ≥ 4/group; two-way ANOVA with Sidak’s post hoc test).

## DISCUSSION

Gemcitabine resistance remains a central barrier to improving outcomes in PDAC ^60,61^. Herein, we identify INPP4B-driven lysosomal exocytosis as a mechanism that lowers intracellular gemcitabine retention, reduces drug-induced DNA damage, and promotes tumour-cell survival. INPP4B expression correlated with gemcitabine resistance and increased cell-surface LAMP1, a functional marker of lysosomal exocytosis. Gain- and loss-of-function experiments further demonstrate that Inpp4b functionally drives this resistance through a PIKfyve–TRPML1 lysosomal trafficking axis ^25^.

Our findings extend the concept of lysosome-mediated drug resistance beyond the classical sequestration of lipophilic weak-base agents ^59^. Gemcitabine is a hydrophilic nucleoside analogue whose activity requires intracellular uptake and metabolic activation before incorporation into DNA ^12^. LC–MS/MS analyses showed increased extracellular gemcitabine following treatment of Inpp4b-overexpressing cells and inhibition of lysosomal function, PIKfyve, or TRPML1 reduced extracellular gemcitabine release and reduced γH2AX levels. These data support a model in which Inpp4b-dependent lysosomal exocytosis reduces the intracellular availability of gemcitabine and thereby limits its effects. The translational relevance of lysosomal targeting in PDAC is supported by clinical studies of CQ and hydroxychloroquine (HCQ) ^62,63^. In the metastatic setting, addition of HCQ to gemcitabine/nab-paclitaxel increased response rates but did not improve 12-month overall survival in an unselected patient population ^62^. Similarly, neoadjuvant gemcitabine/nab-paclitaxel plus HCQ improved pathological response and pharmacodynamic evidence of autophagy inhibition without a clear survival advantage ^63^. These results indicate that broad lysosomal/autophagy inhibition may enhance chemotherapy activity in some PDACs, but that treatment benefit is likely diluted when patients are enrolled without molecular selection. Our results offer a mechanistic explanation for this clinical heterogeneity and support a potential biomarker-driven strategy for patient selection. We propose that INPP4B-high tumours, especially those with elevated cell-surface LAMP1, represent a PDAC subtype where lysosomal exocytosis is the key driver of resistance. In these cases, lysosomal inhibition is a candidate therapy likely to restore chemotherapy retention, a hypothesis supported by our *in vivo* findings showing that chloroquine selectively sensitized Inpp4b-overexpressing tumours to gemcitabine. These data support the idea that lysosomal inhibitors may be effective as targeted chemosensitizers rather than broad cytotoxic agents, in some settings.

Our findings also suggest that more selective approaches may be required. CQ and HCQ are clinically accessible but broadly affect lysosomal acidification and autophagy, and their variable clinical performance may reflect incomplete pathway inhibition, dose-limiting toxicity, or effects on lysosomal processes unrelated to drug exocytosis ^64^. In our models, inhibitors targeting PIKfyve or TRPML1 also suppressed the INPP4B-associated phenotype, and apilimod showed particularly robust synergy with gemcitabine. These data suggest that the INPP4B–PIKfyve–TRPML1 axis may be a more mechanistically defined therapeutic target ^25^. Although direct clinical evaluation of PIKfyve or TRPML1 inhibition in PDAC remains limited, selective inhibition of lysosomal trafficking and exocytosis may provide a higher therapeutic index than generalized disruption of lysosomal acidification.

Notably our work demonstrated that INPP4B-mediated lysosomal exocytosis may also contribute to multidrug resistance. Inpp4b overexpression in KPC cells reduced sensitivity not only to gemcitabine, but also to irinotecan, oxaliplatin, paclitaxel, and daunorubicin; conversely, INPP4B depletion sensitized cells to each agent. This finding has clinical significance because irinotecan and oxaliplatin are core components of FOLFIRINOX, and paclitaxel is used with gemcitabine in standard PDAC treatment ^65,66^. These findings suggest that INPP4B expression monitoring could identify tumours at risk of cross-resistance to both major frontline regimens. This finding not only supports early biomarker assessment, but also incorporation of lysosomal inhibitors before multidrug-resistant clones become dominant.

We acknowledge some limitations of our work. Our in vivo studies used subcutaneous CDX models that do not fully represent the desmoplastic, hypovascular, immunosuppressive microenvironment of PDAC, which may influence drug delivery, lysosomal activity, and response to lysosome-targeted combinations. Thus further validation in orthotopic, syngeneic, genetically engineered, and patient-derived models is needed, ideally using clinically relevant gemcitabine/nab-paclitaxel and FOLFIRINOX regimens. Further work should define how lysosomal exocytosis alters active intracellular gemcitabine pools, establish the molecular requirement for PIKfyve, TRPML1, and lysosomal fusion machinery using genetic approaches, and determine whether INPP4B expression predicts response in clinical specimens from completed HCQ-based PDAC trials.

In summary, our study points to INPP4B as a driver of gemcitabine resistance in PDAC through lysosomal exocytosis. By demonstrating that lysosomal pathway inhibition restores intracellular drug activity, DNA damage, and tumour progression in Inpp4b-high models, our findings provide a rationale for biomarker-guided chemotherapy–lysosome inhibitor combinations. Prior CQ/HCQ trials suggest that lysosomal inhibition is feasible but insufficiently effective in unselected PDAC; our work now provides a mechanistic basis to identify the patients most likely to benefit and supports development of more selective inhibitors of lysosomal trafficking.

## RESOURCE AVAILABILITY

### Lead Contact

Leonardo Salmena

### Materials availability

All materials presented in this study can be provided by the lead contact upon request.

### Data and Code Availability

This study did not generate new code; any additional data or details necessary to reanalyze the reported findings can be provided by the lead contact upon request.

## ACKNOWLEDGEMENTS

We thank all past and present members of the Salmena Lab for contributions. L.S. is a recipient of Tier II Canada Research Chair and supported through the Human Frontier Career Development Program award. Funding for this research was provided in part by Temerty Faculty of Medicine and Department of Pharmacology and Toxicology, University of Toronto, and awards received from Canada Foundation for Innovation (grant no.: 33505); Cancer Research Society (grant no.: 24261), and the Canadian Institute of Health Research (grant nos.: 487599 and 518878).

## AUTHOR CONTRIBUTIONS

C.M.P.M: Conceptualization, Data curation, Formal analysis, Investigation, Methodology, Project administration, Validation, Visualization, Writing—original draft, Writing—review & editing; C.N: Data curation, Investigation, Methodology; G.S: Investigation, Resources. N.N: Investigation, Methodology; C.Y: Investigation, Methodology; C.W: Resources, Investigation; L.T: Conceptualization, Formal analysis, Visualization; J.T.C: Data curation, Formal analysis, Writing—review & editing; L.S: Conceptualization, Data curation, Formal analysis, Methodology, Project administration, Validation, Visualization, Writing— original draft, Writing—review & editing.

## DECLARATION OF GENERATIVE AI AND AI-ASSISTED TECHNOLOGIES IN THE WRITING PROCESS

During the preparation of this work, the author(s) used artificial intelligence (AI) solely for word processing and spell-checking purposes. After using these tools, the author(s) reviewed and edited the content as needed and take(s) full responsibility for the content of the publication.

## SUPPLEMENTAL INFORMATION

### Supplemental Figures

**Figure S1** Datesets from the CCLE and corresponding AUC responses to gemcitabine-based regimens in the CTRP for, **(a)** gemcitabine combined with tanespimycin and **(b)** gemcitabine combined with navitoclax. **(d)** Pearson correlation analysis between basal INPP4B protein expression and gemcitabine IC50 values was used to determine significance. Significance: \**p* < 0.05, \*\**p* < 0.01, \*\*\**p* < 0.001, \*\*\*\**p* < 0.0001; ns, not significant. All replicate experiments were performed at least three times. Whole-cell lysates from **(c)** PANC-1 and **(e)** PK1 pancreatic cancer cell lines transduced with pSMAL-Puro-Empty or pSMAL-Puro-FLAG-INPP4B lentivirus were subjected to Western blotting analysis for INPP4B **(g)** PK8 Cas9-containing cell lines were transduced with INPP4B-targeting sgRNA constructs (Doench3), including a non-targeting sgRNA control (sgLacZ), and GAPDH (as a loading control). Alamar Blue cell viability assay following 96-hour gemcitabine treatment of **(d)** PANC-1 and **(f)** PK1 cells transduced with pSMAL-Puro-Empty (Empty) or pSMAL-Puro-Flag-INPP4B (INPP4B) lentivirus. **(h)** Alamar Blue cell viability assay following 96-hour gemcitabine treatment of PK8 sgINPP4B cells, with respective sgLacZ controls. P-values were calculated using the two-tailed Student’s t-test to analyze the differences for statistical significance between two groups with the assumption of normal distribution of data and equal sample variance. P-values for gemcitabine dose-response curves were calculated in GraphPad Prism using the two-way ANOVA and Sidak’s test for multiple comparisons.

**Figure S2 (a)** Cell surface LAMP1 fold change in MIAPACA2 pSMAL and Inpp4b overexpression cells. **(b)** Cell surface LAMP1 fold change in MIAPACA2 cells following chloroquine (CQ) or apilimod treatment. **(c)** Cell surface LAMP1 fold change in MIAPACA2 cells following MLSI3 or bafilomycin A1 (Baf) treatment. Data represent mean ± SEM from at least three independent experiments. Significance: \**p* < 0.05, \*\**p* < 0.01, \*\*\**p* < 0.001, \*\*\*\**p* < 0.0001; ns, not significant. Statistical significance was determined by two-way ANOVA followed by Sidak’s post hoc test for multiple comparisons for b-c, and unpaired two-tailed parametric t-test for a.

**Figure S3** Representative DR-curve following **(a)** chloroquine, **(c)** bafilomycin A1, **(e)** apilimod and **(g)** MLSI3 following 72-hour treatment of KPC Inpp4b overexpression cells, with pSMAL control. Calculated IC50 values (µM) for **(b)** chloroquine, **(d)** bafilomycin A1, **(f)** apilimod and **(h)** MLSI3 each cell line. Significance: \**p* < 0.05, \*\**p* < 0.01, \*\*\**p* < 0.001, \*\*\*\**p* < 0.0001; ns, not significant. Data represent mean ± SEM from at least three independent experiments. IC50 values, as calculated on GraphPad Prism. Statistical significance was determined by unpaired two-tailed parametric t-test.

**Figure S4** KPC Inpp4b overexpression cell viability of complete synergy matrix from dose combinations with gemcitabine holding each a single lysosome drug dose constant for **(a)** CQ (n=9), **(b)** Baf (n=9), **(c)** apilimod (n=8) and **(d)** ML-SI3 (n=9). Significance: \**p* < 0.05, \*\**p* < 0.01, \*\*\**p* < 0.001, \*\*\*\**p* < 0.0001; ns, not significant. All replicate experiments were performed at least eight times. Statistical significance was determined by two-way ANOVA followed by Sidak’s post hoc test for multiple comparisons.

**Figure S5** KPC Inpp4b overexpression **(a-b)** combination 6X10 matrix heat maps of synergistic doses of CQ with gemcitabine. CQ combination with gemcitabine **(c)** overall matrix ZIP score and **(d)** Inpp4b overexpression dose combination peak ZIP score compared to control (n=9). **(e)** Cell viability from CQ combination with gemcitabine that achieved peak ZIP score compared to control. **(f-g)** Combination 6X10 matrix heat maps of synergistic doses of Baf with gemcitabine. Baf combination with gemcitabine **(h)** overall matrix ZIP score and **(i)** Inpp4b overexpression dose combination peak ZIP score compared to control (n=9). **(j)** Cell viability from Baf combination with gemcitabine that achieved peak ZIP score compared to control. **(k-l)** combination 6X10 matrix heat maps of synergistic doses of apilimod with gemcitabine. Apilimod combination with gemcitabine **(m)** overall matrix ZIP score and **(n)** Inpp4b overexpression dose combination peak ZIP score compared to control (n=9). **(o)** Cell viability from apilimod combination with gemcitabine that achieved peak ZIP score compared to control. **(p-q)** combination 6X10 matrix heat maps of synergistic doses of ML-SI3 with gemcitabine. ML-SI3 combination with gemcitabine **(r)** overall matrix ZIP score and **(s)** Inpp4b overexpression dose combination peak ZIP score compared to control (n=9). **(t)** Cell viability from ML-SI3 combination with gemcitabine that achieved peak ZIP score compared to control. Significance: \**p* < 0.05, \*\**p* < 0.01, \*\*\**p* < 0.001, \*\*\*\**p* < 0.0001; ns, not significant. All replicate experiments were performed at least three times. Statistical significance was determined by unpaired two-tailed parametric t-test for all.

**Figure S6** Relative growth validation of peak ZIP score combination treatment following 6 day incubation of gemcitabine in KPC control and KPC Inpp4b overexpressing cells with **(a)** CQ, **(b)** Baf, **(c)** apilimod and **(d)** MLSI3 (n=3). Significance: \**p* < 0.05, \*\**p* < 0.01, \*\*\**p* < 0.001, \*\*\*\**p* < 0.0001; ns, not significant. Data represent mean ± SEM from at least three independent experiments. Statistical significance was determined by two-way ANOVA followed by Sidak’s post hoc test for multiple comparisons.

**Figure S7** Flow cytometry gating of KPC control cells following cell permeabilization protocol.

**Figure S8** Extracted Ion Chromatogram, MS1 spectrum for the molecule and its MS2 (fragmentation) spectra (left to right).

**Figure S9 (a)** Representative irinotecan DR-curves in KPC pSMAL and Inpp4b overexpressing cell lines and **(b)** calculated irinotecan IC50 values. **(c)** Representative irinotecan DR-curves in Inpp4b knockdown (Sh36 and Sh38) KPC cell lines and **(d)** calculated irinotecan IC50 values. **(e)** Representative oxaliplatin DR-curves in Inpp4b overexpression KPC cell lines and **(f)** calculated oxaliplatin IC50 values for each cell line. **(g)** Representative oxaliplatin DR-curves in Inpp4b knockdown (Sh36 and Sh38) KPC cell lines and **(h)** calculated oxaliplatin IC50 values for each cell line. **(i)** Representative paclitaxel DR-curves in Inpp4b overexpression KPC cell lines and **(j)** calculated paclitaxel IC50 values for each cell line. **(k)** Representative paclitaxel DR-curves in Inpp4b knockdown (Sh36 and Sh38) KPC cell lines and **(l)** calculated paclitaxel IC50 values for each cell line. **(m)** Representative daunorubicin (DNR) DR-curves in Inpp4b overexpression KPC cell lines and **(n)** calculated DNR IC50 values for each cell line. **(o)** Representative DNR DR-curves in Inpp4b knockdown (Sh36 and Sh38) KPC cell lines and **(p)** calculated DNR IC50 values for each cell line. Significance: \**p* < 0.05, \*\**p* < 0.01, \*\*\**p* < 0.001, \*\*\*\**p* < 0.0001; ns, not significant.All replicate experiments were performed at least three times. IC50 values, as calculated on GraphPad Prism. Statistical significance was determined by two-way ANOVA followed by Sidak’s post hoc test for multiple comparisons for d, h, i and p, and unpaired two-tailed parametric t-test for b, f, j and n.

**Figure S10** Quantification of harvested tumor weights (g) in **(a)** KPC pSMAL xenografts and **(b)** KPC Inpp4b xenografts. **(c)** Calculated inhibition of tumor growth rate (ITR, %) for GEM-treated mice with and without concurrent CQ treatment. Significance: \**p* < 0.05, \*\**p* < 0.01, \*\*\**p* < 0.001, \*\*\*\**p* < 0.0001; ns, not significant.Experimental groups contained a minimum of four mice per condition. IC50 values, as calculated on GraphPad Prism. Statistical significance was determined by two-way ANOVA followed by Sidak’s post hoc test for multiple comparisons.

## KEY RESOURCES TABLE

| REAGENT OR RESOURCE | SOURCE | IDENTIFIER |
| --- | --- | --- |
| **Antibodies** |  |  |
| Gapdh | Cell signalling technology | 97166S |
| β-actin | Cell signalling technology | 4967S |
| CD107a (LAMP1) | Invitrogen | # 50-1071-82 |
| Gamma-H2AX phosphorylation kit | Millipore | #17-344 |
| **Chemicals, peptides, and recombinant proteins** |  |  |
| DMEM | Wisent | 319-005 |
| FBS | Wisent | 080-150 |
| penicillin-streptomycin | Wisent | 450-201 |
| Gemcitabine (*In vitro*) | Selleck | 122111-03-9 |
| Bafilomycin A1 | Selleck | S1413 |
| Apilimod | Selleck | S6414 |
| MLSI3 | Selleck | E0026 |
| Chloroquine | Selleck | S6999 |
| Gemcitabine HCL (*In vivo*) | Sigma | G6423 |
| Poly(2-hydroxyethyl methacrylate) | Sigma | 192066 |
| Puromycin | Sigma | P9620 |
| Growth Factor Reduced Matrigel® Basement Membrane Matrix | Corning | 354230 |
| Protease and phosphatase inhibitors | Thermo Fisher Scientific | PI78441 |
| Protamine sulfate | Sigma | P3369 |
| **Critical commercial assays** |  |  |
| Gamma-H2AX phosphorylation kit | Millipore | #17-344 |
| **Experimental models: Cell lines** |  |  |
| KPC | Cancertools | 153474 |
| MiaPaCa-2 | ATCC | CRL-1420 |
| HPAC | ATCC | CRL-2119 |
| PK-1 | provided by Dr. D. Hedley | Ontario Cancer Institute,<br>Toronto, ON, Canada |
| PK-8 | provided by Dr. D. Hedley | Ontario Cancer Institute,<br>Toronto, ON, Canada |
| PANC-1 | ATCC | CRL-1469 |
| BxPC3 | ATCC | CRL-1687 |
| CAPAN1 | ATCC | HTB-79 |
| CAPAN2 | ATCC | HTB-80 |
| NeuroMab Hybridoma Cell Line N171/17 | UC Davis NeuroMab<br>Hybridomas | 066142-UCD |
| **Experimental models: Organisms/strains** |  |  |
| Nu/Nu Mice (Crl:NU-Foxn1nu) | Charles River | 088 |
| **Oligonucleotides** |  |  |
| N/A |  |  |
| **Recombinant DNA** |  |  |
| Sh36 (ULTRA-3254036-pZIP-mCMV-turboRFP-Puro) | transOMIC technologies inc. | 3254036 |
| Sh38 (ULTRA-3254038-pZIP-mCMV-turboRFP-Puro) | transOMIC technologies inc. | 3254038 |
| NTC (TLNSU4420R-ULTRA-NT#4-pZIP-mCMV-turboRFP-Puro) | transOMIC technologies inc. | TLNSU4420R |
| mInpp4b-pTWIST | TWIST Bioscience |  |
| **Software and algorithms** |  |  |
| Graphpad Prism10 | GraphPad Software | www.graphpad.com |
| SynergyFinder 3.0 | Aittokallio Group | synergyfinder.org |
| ImageJ/Fiji |  | https://doi.org/10.1038/nmeth.2019 |
| CytExpert | Beckman Coulter | RRID:SCR_017217 |

## EXPERIMENTAL MODELS

### Cell lines

The KPC mouse pancreatic cancer cell line and MiaPaca2, BxPC-3, CAPAN-2, CAPAN-1, HPAC, PANC-1, PK-1, and PK-8 human pancreatic cancer cell lines were used in this study. All cells were grown in Dulbecco’s Modified Eagle Medium (DMEM; Wisent) supplemented with 10% fetal bovine serum (FBS; Wisent) and 1% penicillin-streptomycin (Wisent) as complete DMEM media. Cells were maintained at 37°C and 5% CO_2_.

### Animal Models

**Species and Strain:** Athymic nude mice (strain: Nu/Nu; Foxn1^nu^/Foxn1^nu^); **Sex:** Female; **Age/Developmental Stage:** 8-10 weeks old at the time of tumor cell inoculation; **Housing and Maintenance:** Mice were housed in specific pathogen-free (SPF) conditions with controlled temperature (20–24°C), humidity (40–60%), and a 12-hour light/dark cycle. Animals had ad libitum access to standard rodent chow and autoclaved water. Cages, bedding, and all materials were sterilized prior to use to maintain pathogen-free status.

## METHODS

### Drug-Gene Correlation Analysis

The relationship between log10 gemcitabine IC50 and normalized INPP4B protein expression was assessed across 8 cell lines (MiaPaca2, BxPC-3, CAPAN-2, CAPAN-1, HPAC, PANC-1, PK-1, and PK-8). Pearson’s correlation coefficient (r) and 95% confidence intervals were calculated using a two-tailed test. Statistical significance was defined as p < 0.05. All analyses were performed in GraphPad Prism, and data are presented as mean ± SEM.

### Lentiviral Transduction and Stable Cell Line Generation

Lentiviral plasmids were amplified in Stbl3™ E. coli and purified using the QIAGEN Plasmid Maxi Kit. Lentiviral particles were generated by co-transfecting HEK 293T cells with the transfer plasmid, psPAX2, and pMD2.G using calcium phosphate. Viral supernatants were harvested at 48 and 72 hours, pooled, and filtered. Pancreatic cancer cells (2.5 × 10⁵/well) were transduced with viral supernatant supplemented with 8 µg/mL protamine sulfate. Forty-eight hours post-transduction, stable lines were selected using 2 µg/mL puromycin (Sigma-Aldrich) until non-transduced controls were eliminated (∼7 days).

### Generation of Gemcitabine-Resistant KPC Lines

Gemcitabine-resistant KPC cells were established by exposure to escalating drug concentrations. Parental KPC cells were cultured in DMEM supplemented with 10% FBS and 1% penicillin-streptomycin at 37°C/5% CO₂. Treatment began at the experimentally determined IC₅₀ of gemcitabine (dose = 0.060µM), with medium refreshed every 48 hours. Upon reaching 70–80% confluency, surviving cells were passaged and given a 24-hour drug-free recovery period. The drug concentration was subsequently doubled (dose = 0.120µM), and cells were cultured until reaching confluency again. A vehicle control (DMSO) group was maintained in parallel.

### Western Blotting

Cells were lysed in NP40-based buffer supplemented with protease and phosphatase inhibitors. Lysates were normalized using a Bradford assay, resolved by SDS-PAGE, and transferred to nitrocellulose. Membranes were blocked in 5% skim milk and incubated overnight at 4°C with primary antibodies (Supplementary Table x). After incubation with HRP-conjugated secondary antibodies, protein bands were detected using Clarity Western ECL (Bio-Rad) or KwikQuant™ Digital-ECL (Kindle Biosciences) and quantified via ImageJ.

### Hybridoma Culture and Antibody Harvest

NeuroMab Hybridoma Cell Line N171/17 was maintained in DMEM supplemented with 10% FBS and 1% penicillin-streptomycin. To harvest the monoclonal Inpp4b N171/17 clone primary antibody, culture supernatant was collected 24 hours after cells reached 100% confluency.

### Cell Viability Assay

Cell viability was assessed using the AlamarBlue™ assay. Pancreatic cancer cells were seeded in 96-well plates (5.0 × 10³ cells/well) and allowed to adhere overnight. Cells were treated with increasing concentrations of gemcitabine (Sigma) diluted in complete DMEM for 72 hours. AlamarBlue™ reagent (Invitrogen) was added according to the manufacturer’s instructions, and fluorescence was measured using a SpectraMax M3 spectrophotometer (excitation: 555 nm, emission: 585 nm). Results were normalized to vehicle controls (0.1% DMSO), with background signal subtracted using cell-free blanks. All conditions were performed in quadruplicate.

### 3D Spheroid Formation and 3D Invasion Assay

To generate non-adherent surfaces, 96-well round-bottom plates were coated with Poly(2-hydroxyethyl methacrylate) (Poly-HEMA) dissolved in 95% ethanol and evaporated under sterile conditions. RFP-expressing cells were seeded at a density of 3.0×10^3 cells/well in 100 µL complete DMEM and centrifuged at 300g for 10 min. Plates were incubated at 37°C in 5% CO₂ for 72-96 hours to allow self-assembly into compact spheroids. Mature spheroids were transferred to pre-chilled 96-well flat-bottom plates and embedded in 50 µL of Growth Factor Reduced Matrigel® Basement Membrane Matrix. Plates were incubated at 37°C for 45–60 min to promote polymerization, after which 100 µL of complete DMEM was added as a chemoattractant. Fluorescence images were captured immediately (T=0) and at 6-days post polymerization (Day 11) using an EVOS Fluorescence Microscope with RFP filters. Spheroid area and circularity were quantified using ImageJ/Fiji software. To account for initial variations, data were normalized to T=0 values. Area fold change was calculated by dividing the area at a given time point (Tx) by the area of the same spheroid at T=0. This value represents the combined effect of cell proliferation and invasive growth. Similarly, circularity fold change was determined by dividing the circularity value at (Tx) by its value at T=0. An increase in this latter value indicates that the spheroid is becoming less circular and more irregular, which is characteristic of invasive cell migration into the surrounding matrix.

Area fold change was calculated as $Area_{Tx} / Area_{T0}$ to measure proliferation and invasion, while circularity fold change was calculated as $Circularity_{Tx} / Circularity_{T0}$ to assess invasive morphology, where deviation from 1.0 indicates irregular, invasive migration.

### Drug Synergy Analysis

Synergistic interactions were evaluated using a 10×6 dose-response matrix. Cells seeded in 96-well plates were treated with ten concentrations of gemcitabine combined with six concentrations of a lysosome inhibitor (Chloroquine, MLSi3, Bafilomycin A1, or Apilimod). Viability was measured after 96 hours via AlamarBlue™ as described above. Experiments were conducted in biological triplicate with technical duplicates, and data were normalized to DMSO controls. Synergy scores were calculated using the Zero-Interaction Potency (ZIP) model via SynergyFinder 3.0. This model integrates Loewe additivity and Bliss independence to generate a delta (∂) score, where ∂ > 5 denotes synergy, ∂ < - 5 indicates antagonism, and scores near zero reflect additive effects.

### Crystal Violet Proliferation Assay

Pancreatic cancer cells were seeded in 24-well plates (2.5 × 10⁴ cells/well) and proliferation was monitored over 6 days. At each time point (starting 24 hours post-seeding), cells were washed with PBS and fixed in 10% neutral buffered formalin (Sigma) for 10 minutes. Plates were stored in PBS at 4°C until the final time point. Fixed cells were stained with 0.1% crystal violet in 20% methanol for 10 minutes, washed three times with distilled water, and air-dried. Dye was solubilized in 10% acetic acid for 30 minutes with shaking, and absorbance was measured at 595 nm using a SpectraMax M3 spectrophotometer. Relative cell growth was calculated by normalizing the absorbance at Day 6 to the absorbance at Day 0.

### Quantification of Surface LAMP1

To assess lysosomal exocytosis, surface LAMP1 (CD107a) was quantified by flow cytometry. Cells were treated for 24 hours with lysosome inhibitors (Chloroquine, MLSi3, apilimod or Bafilomycin A1) or vehicle control. Cells were harvested, washed with cold FACS buffer (PBS/2% FBS/0.05% NaN3), and incubated on ice for 30 minutes with anti-CD107a-eFluor™ 660 (clone eBioH4A3, Invitrogen). Unstained controls were included for each condition. Samples were analyzed on a CytoFLEX flow cytometer (Beckman Coulter) using the APC channel (Ex 638 nm / Em 660/20 nm). Data represent the Mean APC fluorescence intensity of at least 10,000 single cell events, gated by FSC/SSC and doublet exclusion.

### Quantification of *γ*H2AX via Flow Cytometry

To assess DNA damage, *KPC-pSMAL* and *KPC-Inpp4b* cells (5.0 × 10⁵/well) were treated with gemcitabine for 24 hours. Harvested cells were fixed in 2% formaldehyde/PBS for 20 minutes on ice. permeabilization and staining were performed simultaneously for 20 minutes in saponin buffer (0.5% saponin, 10 mM HEPES pH 7.4, 140 mM NaCl, 2.5 mM CaCl₂) containing anti-phospho-Histone H2A.X (Ser139)-FITC or Normal Mouse IgG-FITC isotype control. Cells were washed with 0.1% saponin/PBS, resuspended in PBS, and analyzed by flow cytometry.

### Subcutaneous Xenograft Model

All animal procedures were approved by the DCM Animal Care Committee (Protocol # 20012513) and followed CCAC guidelines. Female athymic Nu/Nu mice (6–8 weeks, Charles River Laboratories) were housed in SPF conditions. Pancreatic cancer cells were resuspended in PBS/Matrigel® (1:1 v/v) and injected subcutaneously into the right flank (1.0 × 10⁶ cells/mouse). Tumor dimensions and body weight were measured every other day. Tumor volume was calculated as V = (Length \times Width^2)/2. When tumors reached ∼100 mm³, mice were randomized into treatment groups.

### *In Vivo* Therapeutic Study

Mice were treated for two weeks with: (1) Gemcitabine (80 mg/kg, i.p., once weekly); (2) Chloroquine (30 mg/kg, oral gavage, 5 days/week); or (3) the combination of the 2 two drugs or (4) corresponding vehicles. Following treatment, mice were monitored for a one-week follow-up period before termination. After the treatment period, tumors were harvested and weighed, rate of tumor inhibition (ITR) of the drug was calculated by 100% × (average tumor weight of the control-average tumor weight of the treated)/ (average tumor weight of the control).

### LC-MS/MS Methods

Metabolites were separated by reverse-phase ultra-high-performance liquid chromatography (UHPLC) using a Thermo Scientific Ultimate 3000 system coupled to a Waters Atlantis Premier BEH C18 AX column (100 × 2.1 mm, 1.7 µm particle size) fitted with a guard column. The column and autosampler were maintained at 40°C and 5°C, respectively, and 10 µL of each sample was injected. Mobile phase A consisted of 0.1% formic acid in water, and mobile phase B consisted of 0.1% formic acid in acetonitrile, delivered at a flow rate of 0.4 mL/min. The gradient program was as follows: 5% B held from 0–1 min, increased linearly to 98% B from 1–7 min, held at 98% B from 7–10 min, returned to 5% B from 10–10.5 min, and held at 5% B from 10.5–15 min for re-equilibration.

Eluted analytes were analyzed on a Thermo Scientific Q Exactive mass spectrometer equipped with a heated electrospray ionization (HESI-II) source operated in positive ion mode. Source parameters were as follows: spray voltage, 3.5 kV; capillary temperature, 320°C; sheath gas, 40 (arbitrary units); auxiliary gas, 20 (arbitrary units); spare gas, 5 (arbitrary units); and S-lens RF level, 55. Full-scan MS data were acquired over an m/z range of 100–350 at a resolution of 70,000 (full width at half maximum), with an automatic gain control (AGC) target of 3 × 10⁶ and a maximum injection time of 200 ms. Data-independent MS2 acquisition targeting the precursor ion at m/z 264.0790 (isolation window, 1.0 m/z) was performed at a resolution of 17,500, with an AGC target of 2 × 10⁵, a maximum injection time of 10 ms, and a normalized collision energy (NCE) of 35. Raw data files were processed using Thermo Scientific Xcalibur Qual Browser software. The analyte of interest was identified and quantified based on the precursor ion at m/z 264.0790 and its corresponding product ion at m/z 112.0508, using a mass accuracy window of 5 ppm. We tested various doses of gemcitabine and identified a 10µM dose was optimal for our analysis. A standard curve was generated against known concentrations of gemcitabine (y = 1.710358E+07x + 7.459938E+05; R² = 9.999370E-01) and used to quantify extracellular concentration.

## QUANTIFICATION AND STATISTICAL ANALYSIS

### Statistical Software and Data Analysis

All statistical analyses were performed using GraphPad Prism (version should be specified) unless otherwise noted. Statistical significance was defined as p < 0.05 (two-tailed tests unless otherwise noted). Where applicable, specific p-value thresholds are denoted as follows: *p < 0.05, **p < 0.01, ***p < 0.001, ****p < 0.0001; ns, not significant. Parametric statistical tests (e.g., t-tests, ANOVA) were applied based on the assumption of normal distribution and homogeneity of variance, which are generally valid for biological replicates with n ≥ 3. For datasets with small sample sizes or evident non-normal distribution, non-parametric alternatives were considered, as indicated in figure legends. The specific statistical test applied to each experiment (e.g., unpaired t-test, one-way ANOVA, two-way ANOVA with post hoc comparisons) is detailed in the corresponding figure legend.

