## Supplementary figures and images for "Inositol Polyphosphate-4-Phosphatase Type II promotes gemcitabine resistance in pancreatic ductal adenocarcinoma cells via lysosomal exocytosis"

### Supplemental 1

#### Melo *et al.* *Supplemental* Figure 1

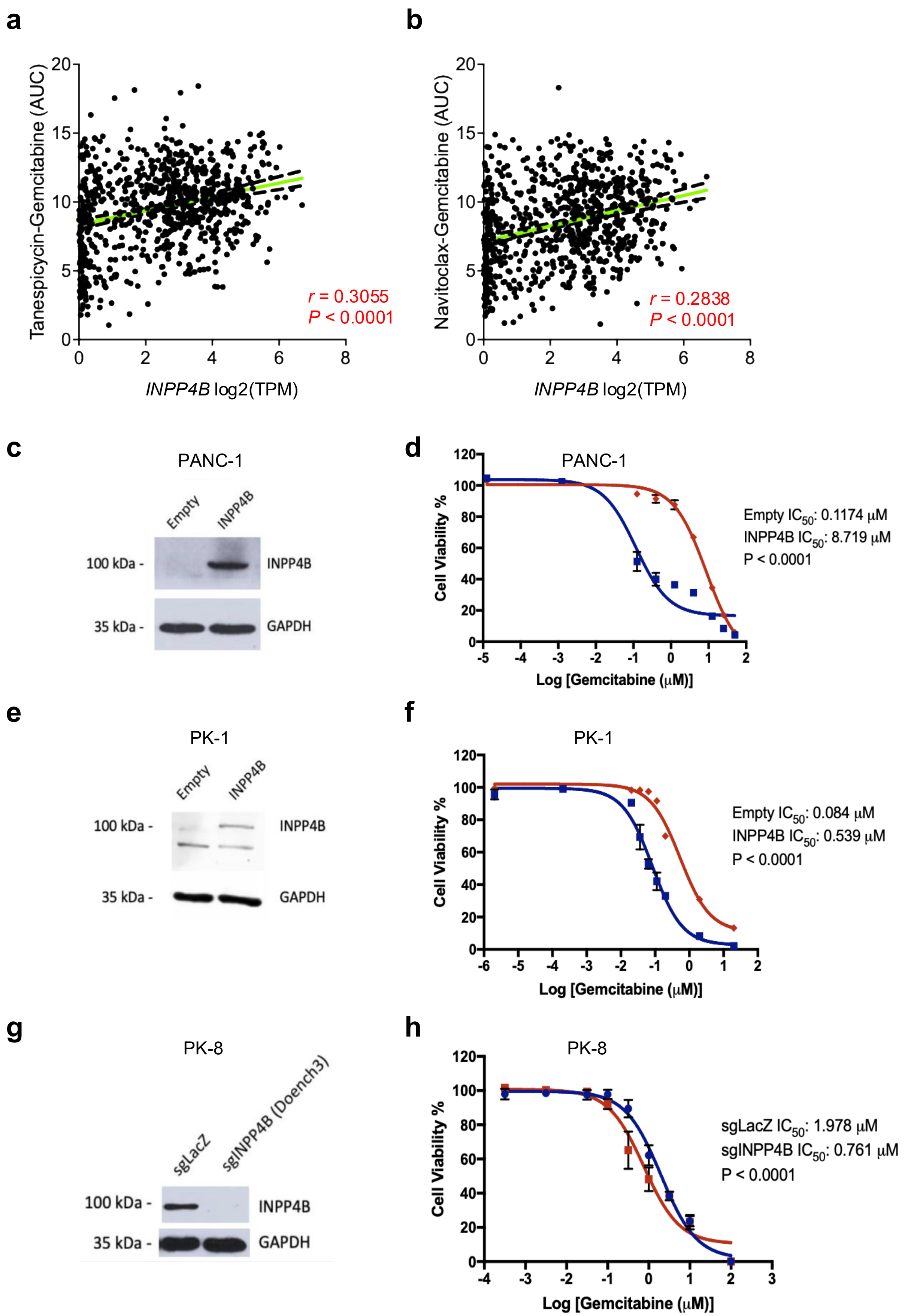

### Supplemental 2

**Melo *et al.* Supplemental Figure 2**

**a**

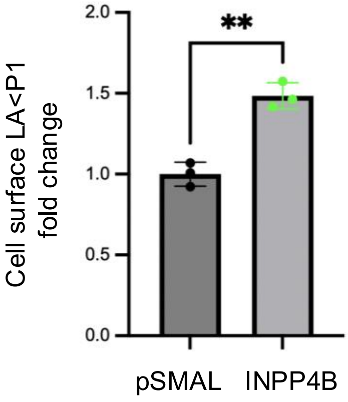


**b**

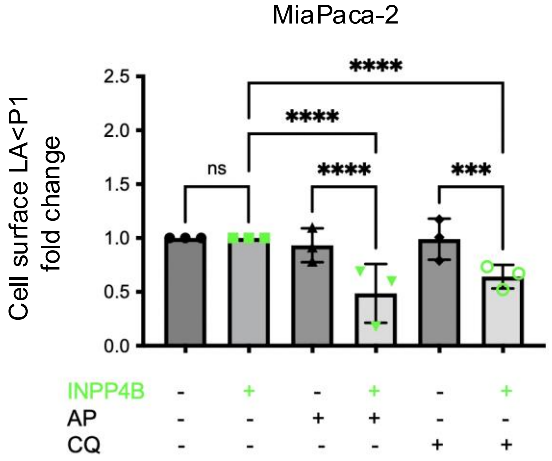


**c**

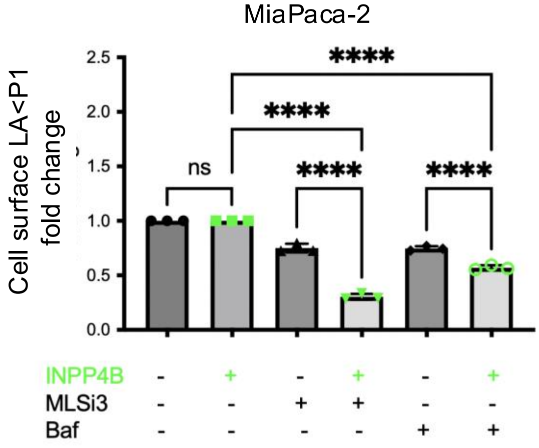

### Supplemental 3

**Melo *et al. Supplemental* Figure 3**

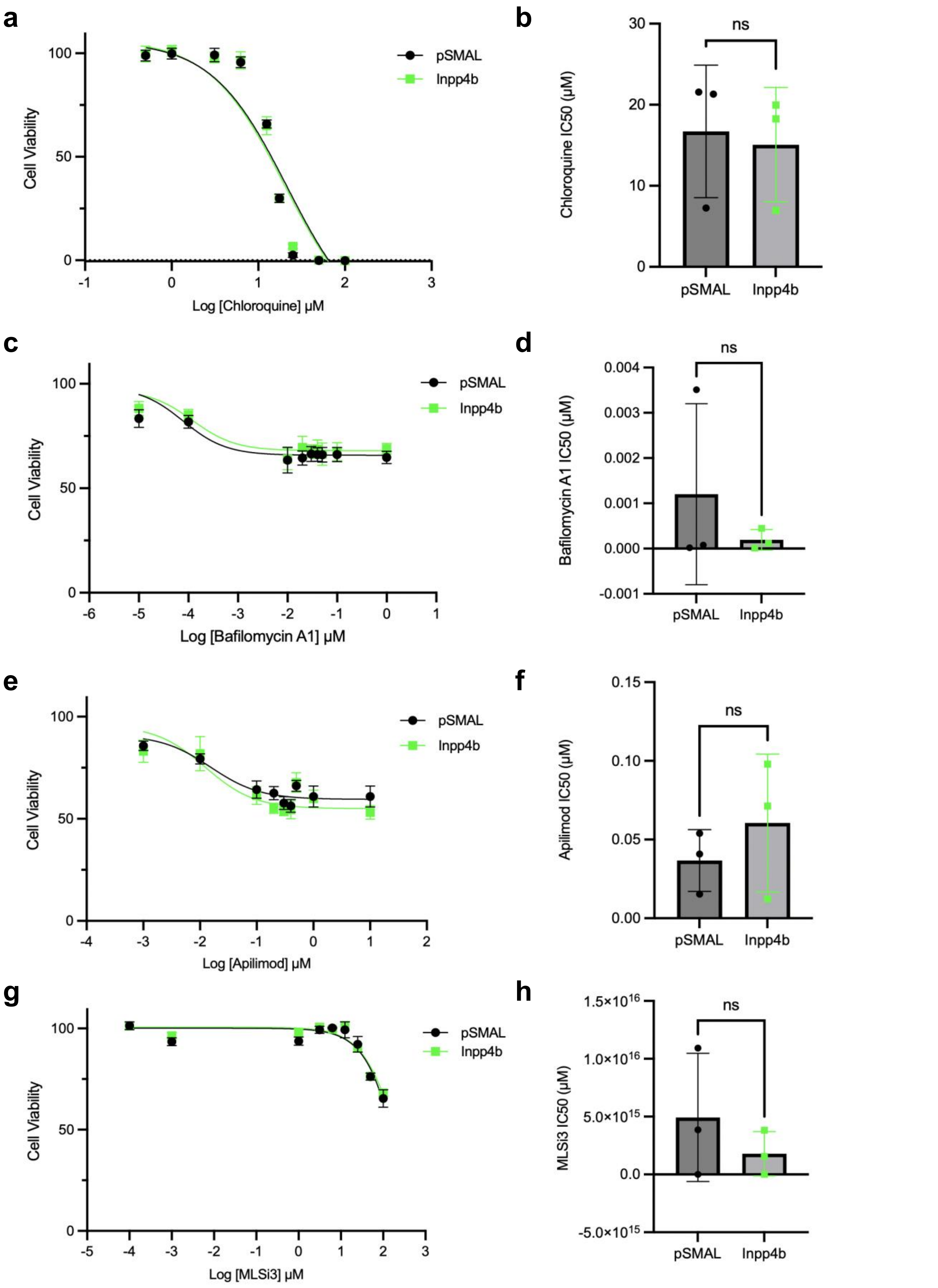

### Supplemental 4

**Melo *et al.* Supplemental Figure 4**

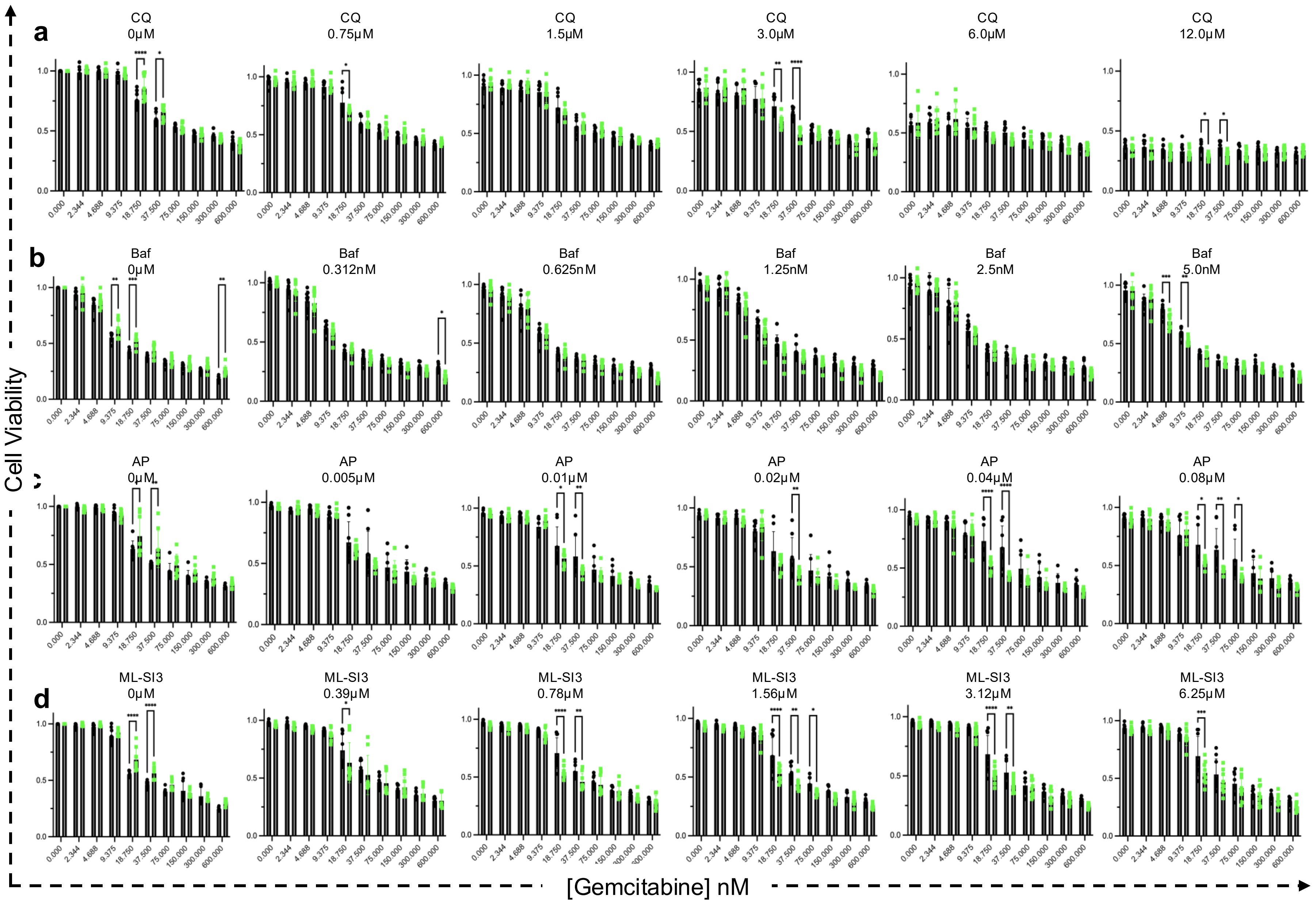

### Supplemental 5

#### Melo *et al.* Supplemental Figure 5

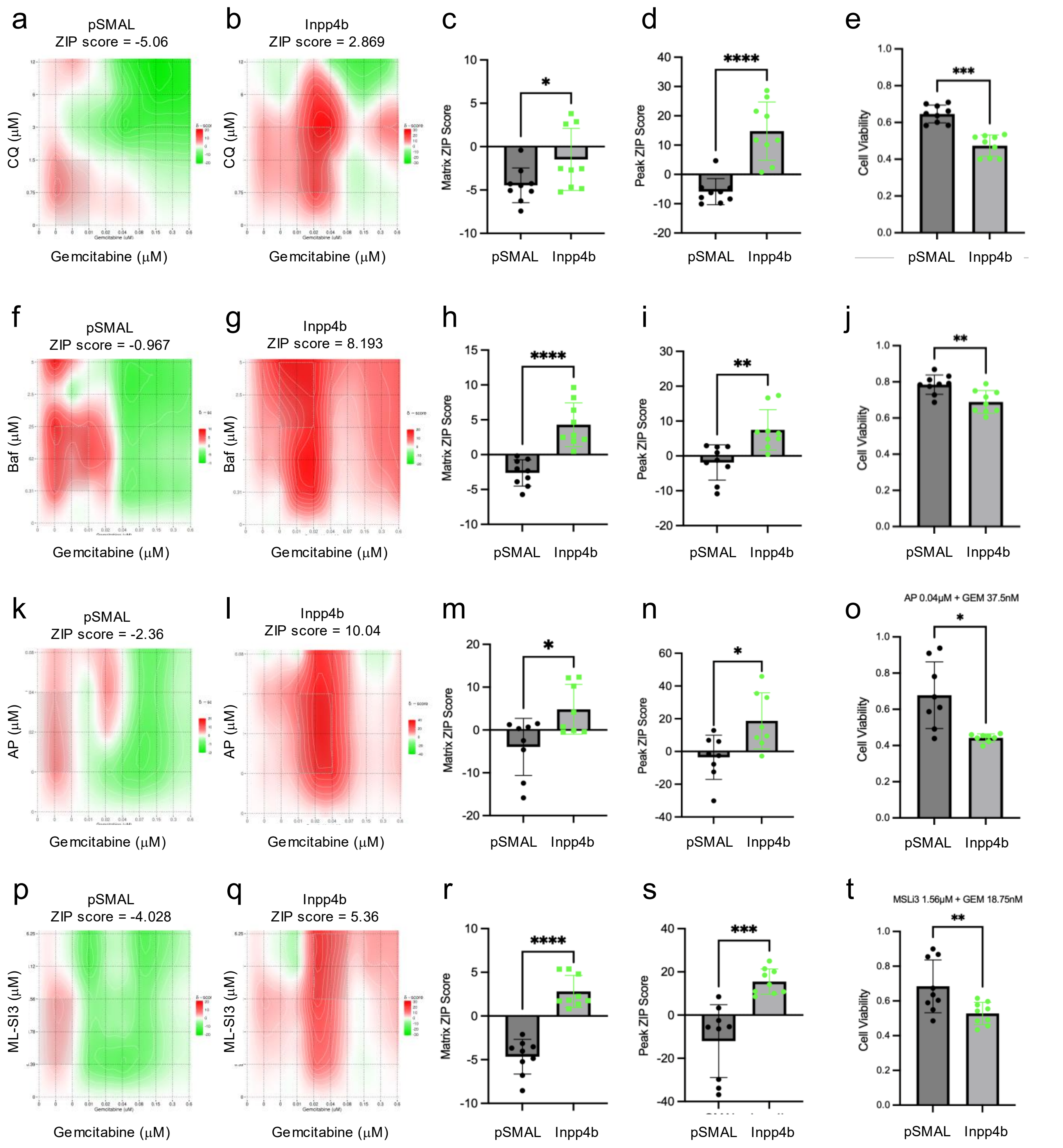

### Supplemental 6

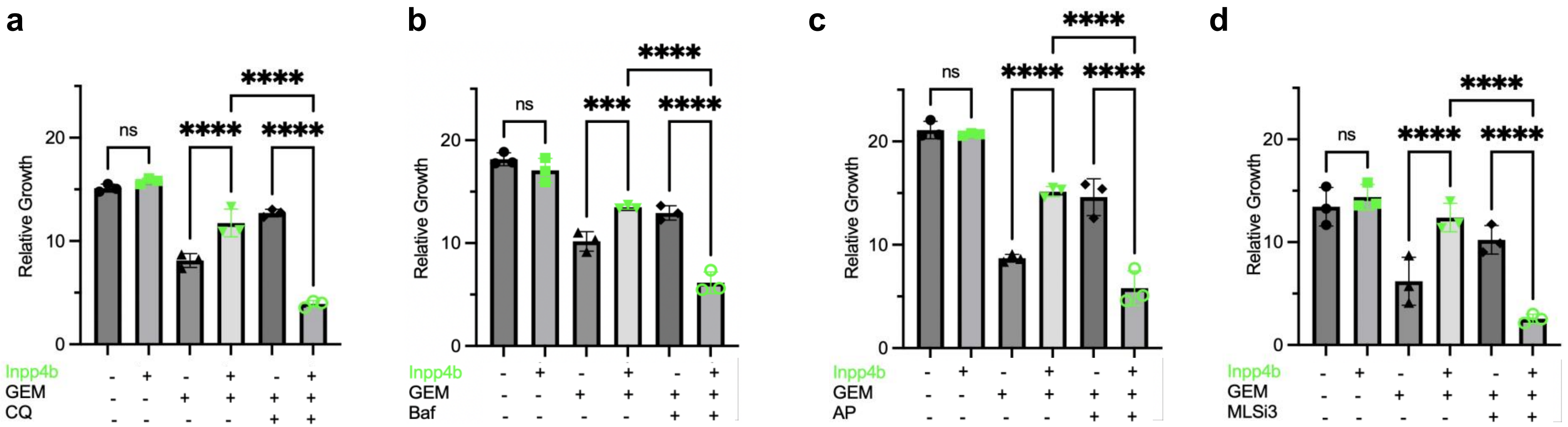

### Supplemental 7

**Melo *et al.* Supplemental Figure 7**

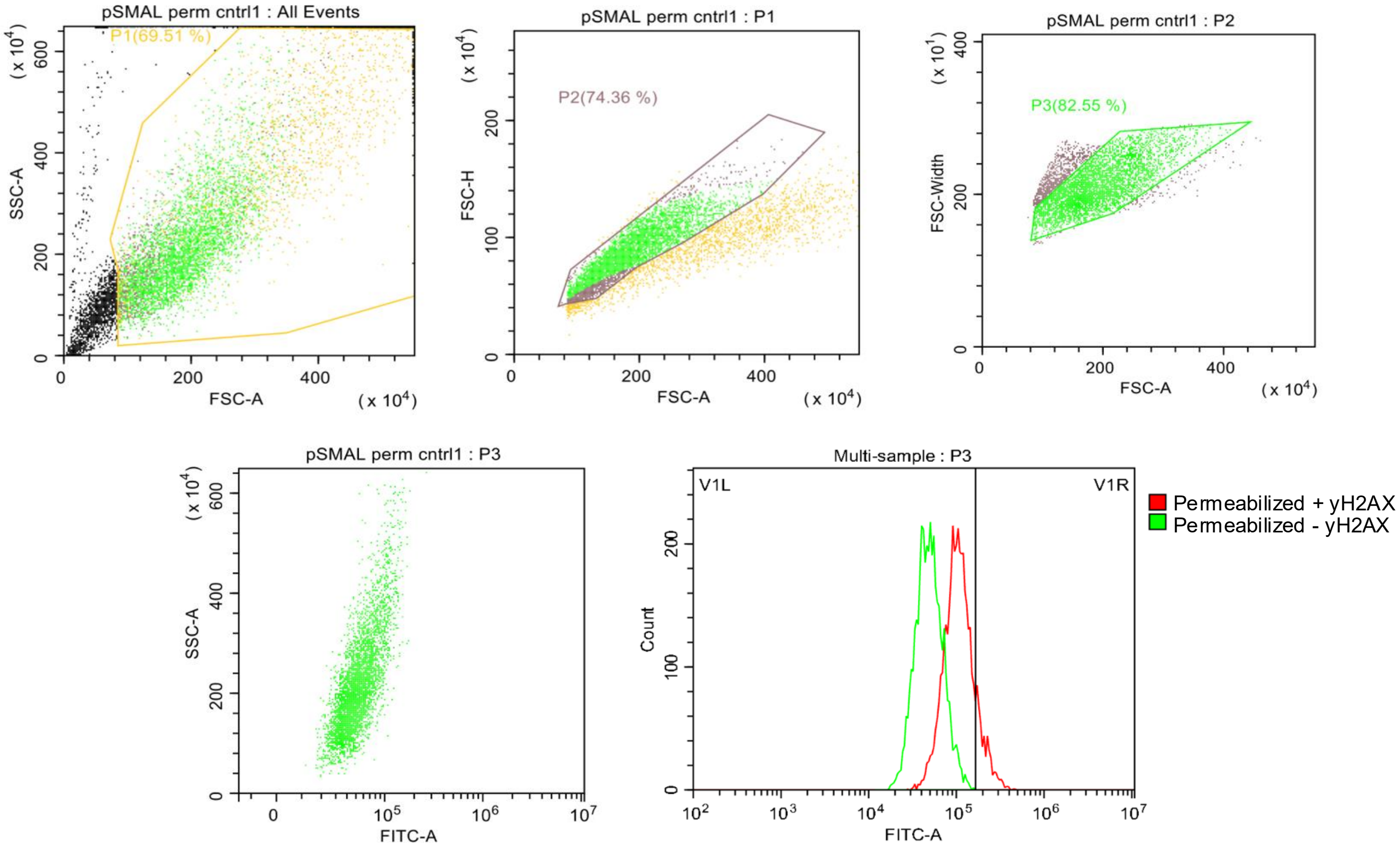

### Supplemental 8

**Melo *et al.* *Supplemental* Figure 8**

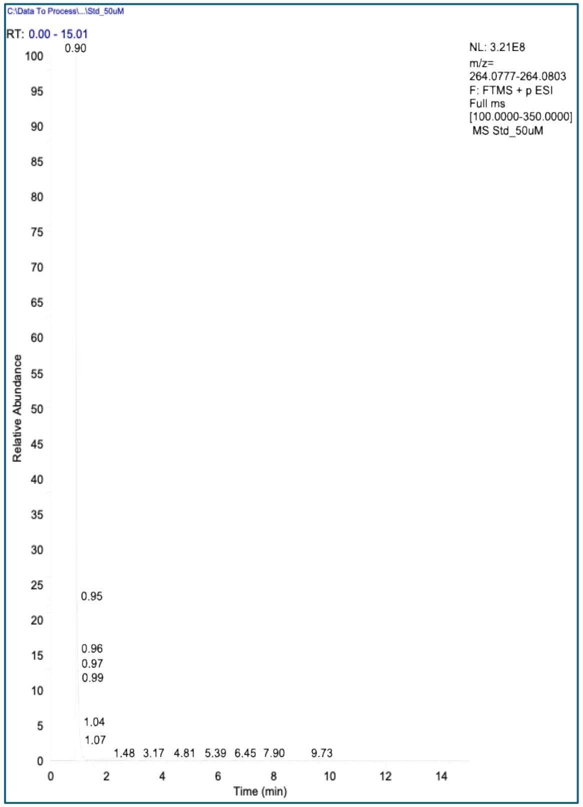


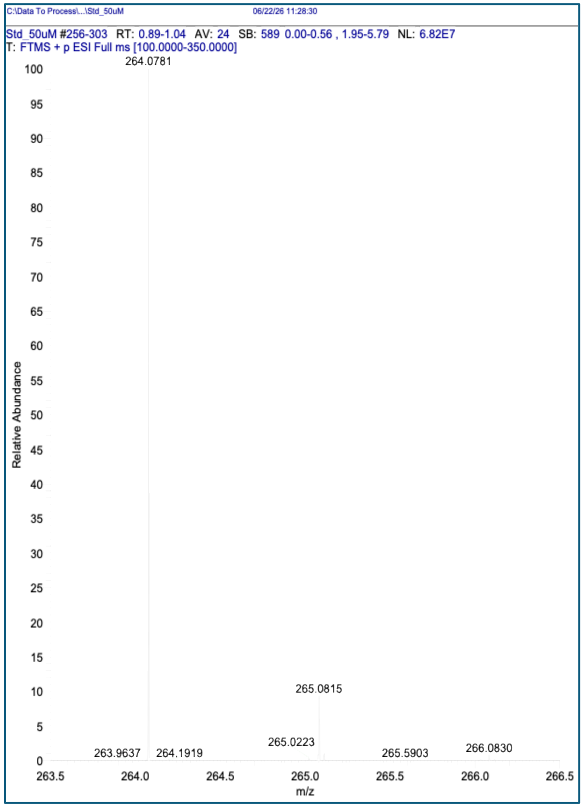


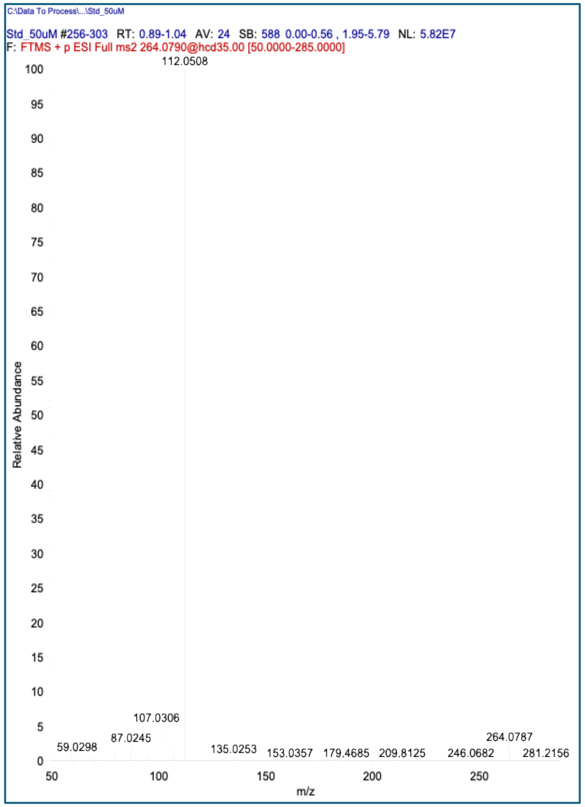

### Supplemental 9

**Melo *et al. Supplemental* Figure 9**

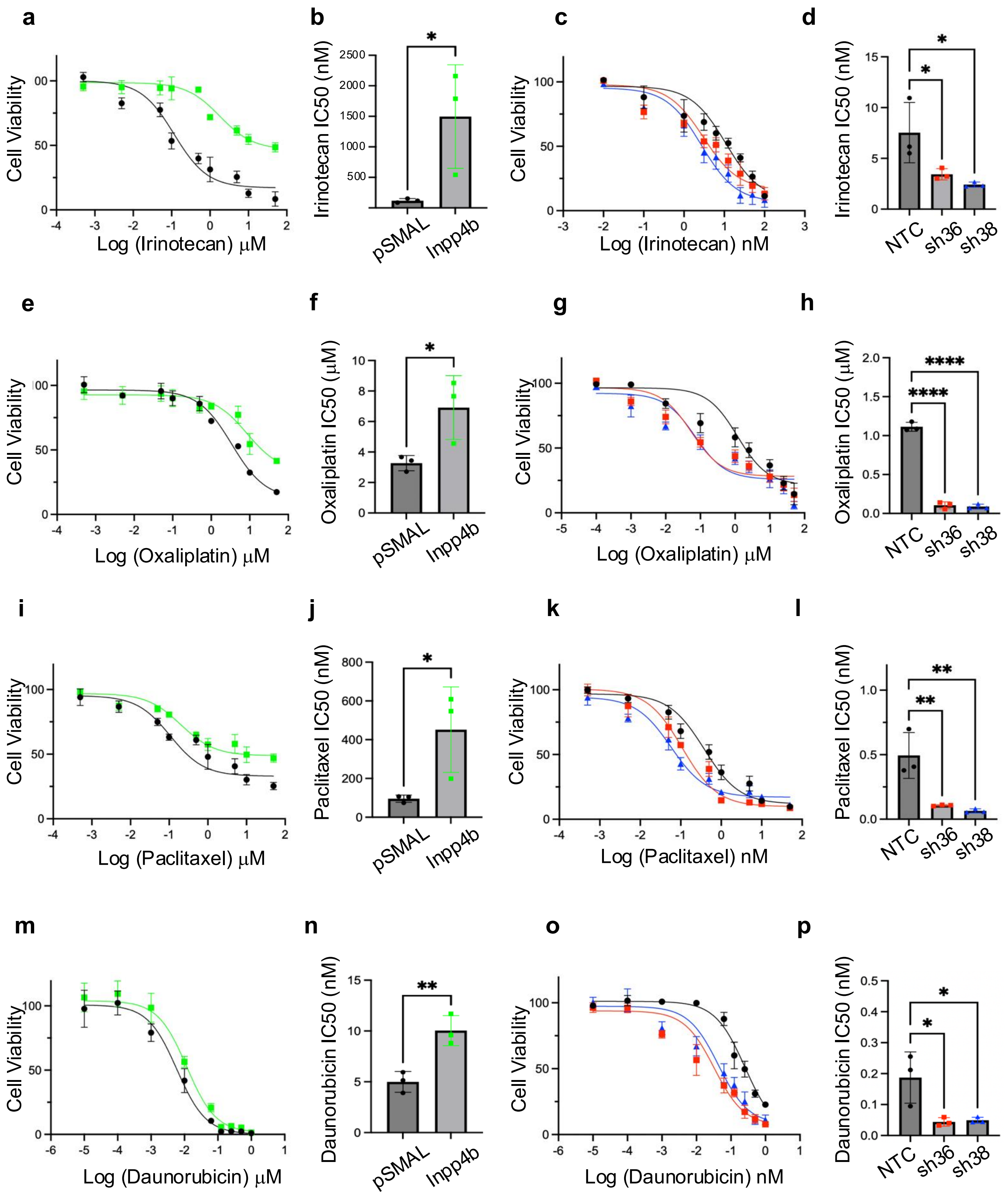

### Supplemental 10

**Melo *et al. Supplemental* Figure 10**

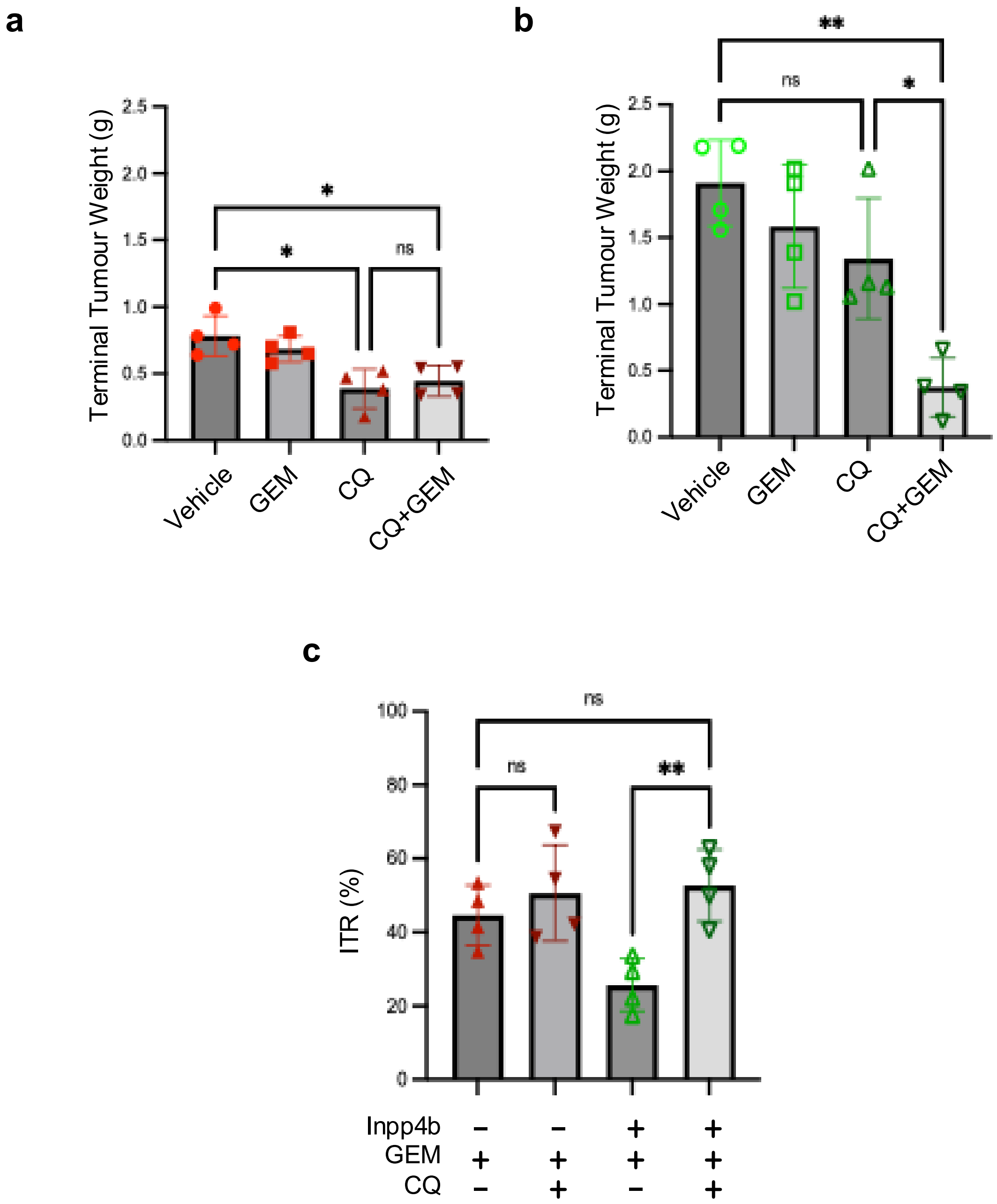
